# SARS-CoV-2 Infects Peripheral and Central Neurons Before Viremia, Facilitated by Neuropilin-1

**DOI:** 10.1101/2022.05.20.492834

**Authors:** Jonathan D. Joyce, Greyson A. Moore, Poorna Goswami, Telvin L. Harrell, Tina M. Taylor, Seth A. Hawks, Jillian C. Green, Mo Jia, Neeharika Yallayi, Emma H. Leslie, Nisha K. Duggal, Christopher K. Thompson, Andrea S. Bertke

## Abstract

Neurological symptoms associated with COVID-19, acute and long-term, suggest SARS-CoV-2 affects both central and peripheral nervous systems. Although studies have shown olfactory and hematogenous entry into the brain and neuroinflammation, little attention has been paid to the susceptibility of the peripheral nervous system to infection or to alternative routes of CNS invasion. We show that neurons in the central and peripheral nervous system are susceptible to productive infection with SARS-CoV-2. Infection of K18-hACE2 mice, wild-type mice, golden Syrian hamsters, and primary neuronal cultures demonstrate viral RNA, protein, and infectious virus in peripheral nervous system neurons and satellite glial cells, spinal cord, and specific brain regions. Moreover, neuropilin-1 facilitates SARS-CoV-2 neuronal infection. Our data show that SARS-CoV-2 rapidly invades and establishes a productive infection in the peripheral and central nervous system via direct invasion of neurons prior to viremia, which may underlie some cognitive and sensory symptoms associated with COVID-19.

## Introduction

Up to 80% of people infected with SARS-CoV-2, the virus responsible for COVID-19, report neurological symptoms. These symptoms span the central nervous system (CNS; anosmia, dizziness, headache, cognitive/memory deficits) and peripheral nervous system (PNS-sensory and autonomic systems; impaired taste/sensation, orthostatic intolerance, syncope)^1–3^. Fatigue, memory issues, “brain fog,” hyper/hypoesthesia, and autonomic dysfunction can persist as part of post-acute sequalae of COVID-19 (PASC; “long COVID”)^4^. Detection of virus, viral RNA, and antigen, in cerebrospinal fluid and brains of COVID-19 patients indicates SARS-CoV-2 is neuroinvasive, which has been documented for common-cold and epidemic coronaviruses (HCoV-OC43, HCoV-229E, MERS, SARS)^5–13^.

As anosmia is a primary symptom of COVID-19, early studies assessed CNS invasion via olfactory sensory neurons (OSNs), using transgenic mice expressing human angiotensin converting enzyme 2 (hACE2), the receptor for SARS-CoV-2^14–16^. Since RNA, protein, and virus were detected in OSNs and sustentacular cells, little attention has since been paid to alternative routes of neural invasion or to the role of PNS infection. PNS symptoms are often reported among non-hospitalized COVID-19 patients, who constitute the bulk of those infected, and up to one-third of those with PASC. Thus, understanding PNS susceptibility to infection and the functional consequences thereof is essential^17, 18^.

Therefore, we assessed the susceptibility of PNS sensory (lumbosacral dorsal root ganglia-LS-DRG, trigeminal ganglia-TG) and autonomic (superior cervical ganglia-SCG) neurons to infection with SARS-CoV-2 following intranasal inoculation of K18-hACE2 transgenic mice (hACE2 mice), wild-type C57BL/6J mice (WT), and golden Syrian hamsters. We also assessed neuroinvasion of the spinal cord and specific brain regions (olfactory bulb, cortex, hippocampus, brainstem, cerebellum), characterized viral growth kinetics in primary neuronal cultures, and investigated the contribution of neuropilin-1 (NRP-1) to neuronal entry.

We show that SARS-CoV-2 productively infects PNS sensory neurons and that autonomic neurons, while susceptible to infection, sustain significant cytopathology and do not release infectious virus. Recovery of infectious virus from the TG, which innervates oronasal mucosa and extends central projections into the brainstem, suggests an alternative route of neuroinvasion independent of OSNs. We also recovered infectious virus from spinal cord and LS-DRGs, which may underlie some sensory disturbances experienced in COVID-19. We further show that SARS-CoV-2 is capable of replicating in specific brain regions, with highest RNA concentrations and viral titers found in the hippocampus, which may contribute to memory disturbances associated with COVID-19. Furthermore, invasion of the brain and PNS occurs rapidly after infection, before viremia, demonstrating direct entry and transport through neurons. Our detection of viral RNA and infectious virus in brains and peripheral ganglia of WT mice and golden Syrian hamsters demonstrates that neuroinvasion can occur independent of hACE2 and is at least partially dependent on NRP-1.

## Results

To assess susceptibility of PNS and CNS neurons to SARS-CoV-2 infection, we intranasally (IN) inoculated hACE2 and WT mice with 10^3^ or 10^5^ PFU SARS-CoV-2 USA-WA1/2020 (n=12/group; Extended Data Fig. 1a, Supplementary Fig. 1a,b). Animals were monitored daily and tissues collected three- and six-days post-infection (dpi). Weight loss began 3 dpi and death occurred 6 dpi (14%) in hACE2 mice inoculated with 10^5^ PFU (Extended Data Fig. 1b). RT-qPCR showed low-level transient viremia and lung infection (Extended Data Fig. 1c). These results are similar to previous studies and demonstrate successful infection^16, 19, 20^.

### PNS sensory TG and sympathetic SCG neurons innervating the oronasopharynx are susceptible to infection

Both sensory and sympathetic pathways through TGs and SCGs, respectively, could serve as neural routes for neuroinvasion. Trigeminal nerves provide sensory innervation to oronasal mucosa, projecting into the brainstem. SCGs provide sympathetic innervation to salivary and lacrimal glands and vasculature of the head and brain, with preganglionic neurons residing in the spinal cord. To assess susceptibility of these peripheral neurons to infection with SARS-CoV-2, TGs and SCGs were assessed for viral RNA, protein, and infectious virus. We detected viral RNA at 3- and 6-dpi in TGs and SCGs in both inoculum groups in hACE2 and WT mice. Viral RNA concentrations were lower in WT than hACE2 mice (Fig. 1a,b), but increased over time in TGs and SCGs of both, suggesting genome replication. Immunostaining detected nucleocapsid (SARS-N) in ≈41% of TG neurons of hACE2 mice and ≈37% of TG neurons of WT mice (Fig. 1c,d, Extended Data Fig. 2a). Nearly all SCG neurons from hACE2 mice (≈97%) were SARS-N-positive, showing substantial pathology with vacuolated neurons and loss of ganglionic architecture. Although SCGs from WT mice remained intact, SARS-N was evident in ≈93% of neurons and some vacuolization was observed. Immunostaining detected spike (SARS-S) in both ganglia similar to detection of SARS-N (Fig. 1e, Extended Data Fig. 2b, antibody validation in Supplementary Fig. 1c).Viral genome replication was confirmed by double-stranded RNA (dsRNA) immunostaining in both ganglia (Fig. 1f, Extended Data Fig. 2c). Infectious virus was recovered in TGs from one 10^5^ PFU-inoculated hACE2 mouse at 3 dpi (5 PFU/mg homogenate) and another at 6 dpi (2 log PFU/mg homogenate). Infectious virus was not detected from SCGs. The pathology of the ganglia and H2AX immunostaining (Supplementary Fig 2) suggest SARS-CoV-2 may have produced such significant cytotoxicity that production of viral progeny was impossible. The use of multiple complementary assays indicate that viral RNA and protein can be isolated from both TGs and SCGs of hACE2 and WT mice, and that infectious virus can be recovered from TGs. Thus, TGs and SCGs are susceptible to infection with SARS-CoV-2, TGs may serve as a route of CNS invasion, and neuroinvasion can occur independent of hACE2.

**Fig 1.**
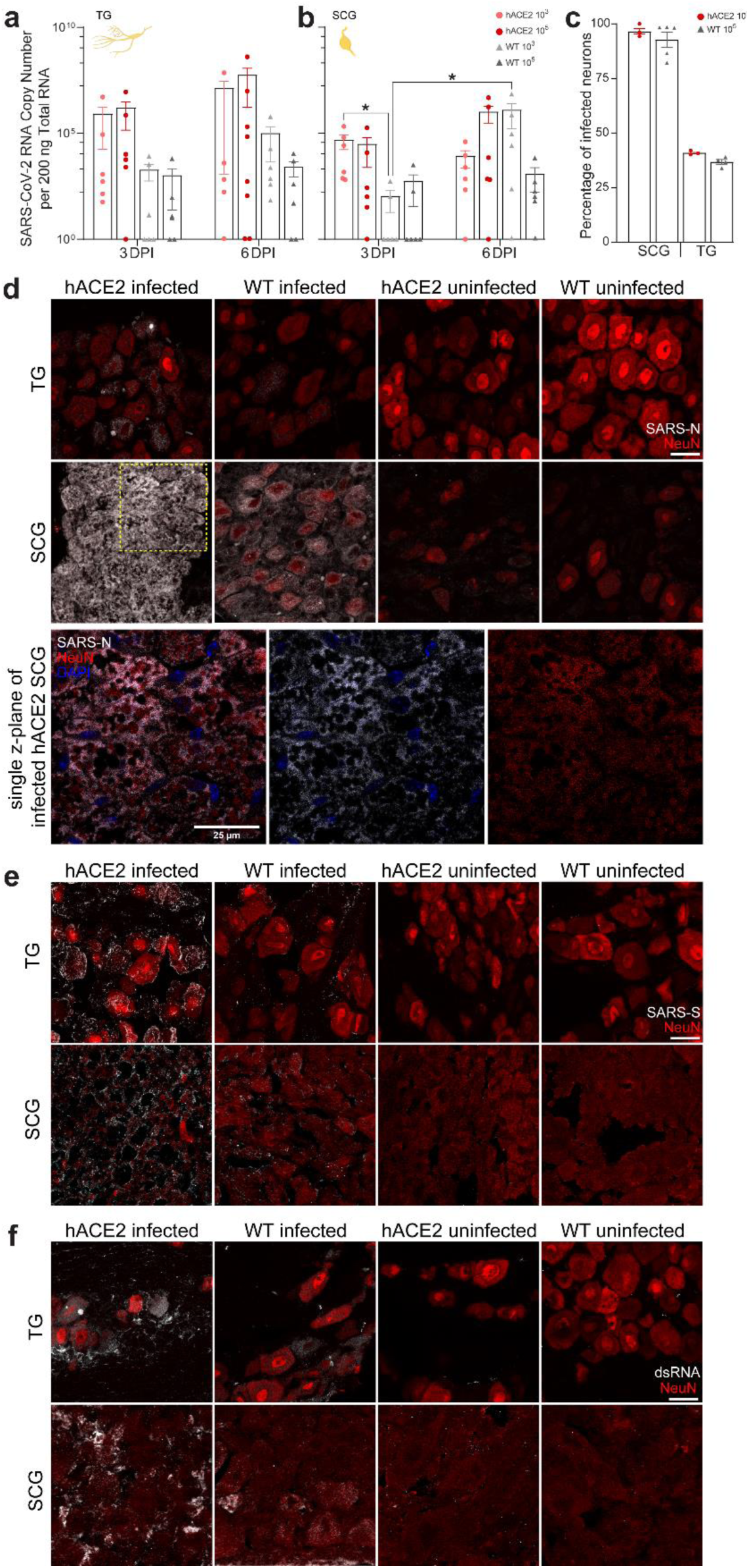
SARS-CoV-2 infection of TG and SCG in hACE2 and wildtype (WT) mice. **a**, SARS-CoV-2 RNA was detected at increasing concentration in TGs of hACE2 and WT mice in both inoculum groups from 3-6 dpi. The TG provides sensory innervation to the face, including the nasal septum, and sends projections to the brain stem, thereby providing an alternative entry point for SARS-CoV-2. No statistically significant differences were detected between the groups (F(7, 40) = 1.855, P = 0.1032). **b**, SARS-CoV-2 RNA was detected at increasing concentration in SCGs of hACE2 and WT mice in both inoculum groups from 3-6 dpi. The SCG provides sympathetic innervation to the salivary glands, lacrimal glands, and blood vessels of the head, neck, and brain. Three-way ANOVA detected a significant difference (F(7, 40) = 3.118, P = 0.0101) in RNA genome copy number. Tukey’s honestly significant difference (HSD) post hoc tests detected significant differences between the WT groups inoculated with 10^3^ PFU assessed at 3- and 6-dpi (*p*=0.048). **c**, Percentage of SCG and TG neurons infected by 6 dpi in tissue sections from 10^5^ PFU-inoculated hACE2 and WT mice. SCGs showed high levels of infection in each mouse type (93-96%) as did TGs (37-41%). **d**, Immunofluorescence for SARS-CoV-2 nucleocapsid (SARS-N, grey) and NeuN (red) labeled neurons in TG and SCG sections at 6 dpi. SARS-N is more prevalent in hACE2 than in WT but observable in both. No SARS-N was detected in ganglia from uninfected animals. Neurons in SCG were particularly sensitive to infection; significant vacuolization was observed in infected hACE2 SCG cells, visible in single z-plane. Contrast for NeuN was increased in the z-plane to better illustrate residual NeuN immunoreactivity inside SARS-N-negative vacuoles in the yellow box above. This cytopathology was common across numerous SCGs in hACE2 mice. **e**, Immunofluorescence for SARS-CoV-2 spike (SARS-S, grey) and NeuN (red) labeled neurons in TG and SCG sections at 6 dpi. Immunostaining was similar to that for SARS-N, with greater SARS-S in hACE2 neurons, but present in both. Vacuolization was again observed in SCG neurons. SARS-S was absent in uninfected neurons. **f**, Immunofluorescence for double stranded RNA (dsRNA, grey) and NeuN (red) labeled neurons in TG and SCG sections at 6 dpi. Immunostaining for dsRNA, a marker of viral replication, was similar to SARS-S and SARS-N immunostaining in TGs and SCGs. dsRNA was present in greater concentrations in SCGs than TGs and in hACE2 mice than WT mice but was present in both ganglia and both mouse types. Positive dsRNA immunostaining indicates SARS-CoV-2 genome replication in TGs and SCGs. Some nonspecific extracellular binding was observed in uninfected mice, but no intracellular immunofluorescence was observed. Data are the mean ± s.e.m. Log transformed RNA genome copy numbers were statistically compared by three-way ANOVA (independent variables: inocula, days post infection, genotype). Pairwise comparisons were conducted using Tukey’s HSD post hoc tests. \**p* < 0.05. n=6 for all animals/timepoints/tissues (except 10^5^ hACE2 6 dpi TG: n=7; 10^3^ hACE2 6 dpi TG: n=5). See Extended Data Figure 2 for unmerged images. See Supplementary Figure 1 for additional antibody validation via western blot, hACE2 genotyping, hACE2 protein expression, and RT-qPCR controls.

### PNS sensory lumbosacral-DRGs and CNS spinal cord neurons are susceptible to infection

Extending our investigation beyond PNS innervations of the oronasopharynx, we assessed the LS-DRG and spinal cord as above. The LS-DRG conveys sensory information (pain, pressure, position) from the periphery and internal organs to the spinal cord. Similar to our results from TGs, we detected viral RNA at 3- and 6-dpi in LS-DRGs in both inoculum groups in hACE2 and WT mice (Fig. 2a). RNA concentrations increased over time, suggesting genome replication. Similar results were found in the spinal cord (Fig. 2b). Immunostaining demonstrated SARS-N in ≈42% of LS-DRG neurons of hACE2 mice and ≈24% of WT neurons (Fig. 2c,d, Extended Data Fig. 2a). Immunostaining for SARS-S in LS-DRGs was similar to that of SARS-N, with SARS-S present throughout hACE2 LS-DRGs, and also in satellite glial cells (SGCs) (Fig. 2e,g Extended Data Fig. 2b, Supplementary Video 1). Immunostaining for dsRNA mirrored that of SARS-N and SARS-S, indicating viral genome replication (Fig. 2f, Extended Data Fig. 2c). Infectious SARS-CoV-2 (2 PFU/mg homogenate) was recovered from LS-DRG homogenate of one hACE2 mouse at 6 dpi, verifying productive infection. Presence of viral RNA and infectious virus in neurons with no axonal targets in the head or lungs suggested spread via hematogenous dissemination or via axonal transport from a distal site of infection. Given that LS-DRGs project to the spinal cord, and transmission from the LS-DRG to the cord (or vice versa) is possible, the spinal cord was also immunostained. We identified punctate SARS-N staining inside spinal cord neurons, as well as diffuse SARS-N signal throughout the spinal cord (Fig. 2h, Extended Data Fig. 3, Supplementary Video 2) suggestive of free virus, or at least N protein, in the cord. Immunostaining for microglia (Iba1, Fig. 2i) and astrocytes (S100b, Fig. 2j) showed minimal co-localization between SARS-N and these cell markers. Infectious virus (4 log PFU/mg homogenate) was recovered from the spinal cord of one hACE2 mouse at 6 dpi, demonstrating productive infection. These data further demonstrate that PNS sensory neurons, as well as neurons in the spinal cord, are susceptible to infection with SARS-CoV-2.

**Fig 2.**
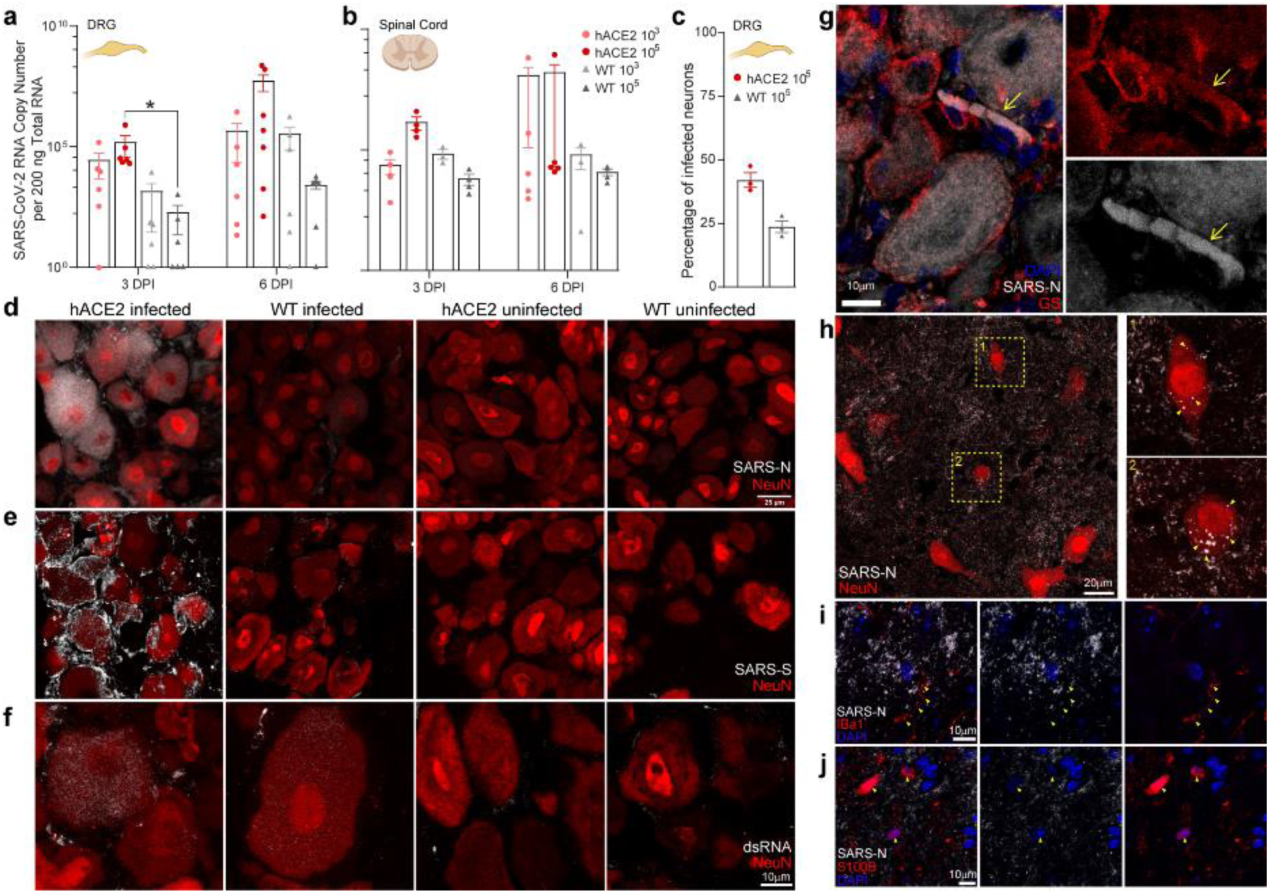
SARS-CoV-2 infection of LS-DRG, including satellite glial cells, and lumbosacral spinal cord in hACE2 and WT mice. **a**, SARS-CoV-2 RNA was detected in increasing concentrations in LS-DRGs of hACE2 and WT mice in both inoculum groups from 3-6 dpi. The LS-DRG conveys sensory information (pain, pressure, position) from the periphery and organs to the spinal cord. Three-way ANOVA detected a significant difference (F(7, 41) = 4.590, P = 0.0007) in RNA genome copy number. Tukey’s HSD detected differences between the hACE2 and WT groups inoculated with 10^5^ PFU at 3 dpi (*p*= 0.018). n=6 for all animals/timepoints (except 10^5^ hACE2 6 dpi: n=7). **b**, SARS-CoV-2 RNA was detected in lumbosacral spinal cords of hACE2 and WT mice in both inoculum groups at both time points. No statistically significant differences were detected between the groups (F(7, 25) = 1.3054, P = 0.2885). Samples sizes were 4 for 10^5^ hACE2 mice at 3 dpi and 5 at 6 dpi; 5 for all timepoints for 10^3^ hACE2 mice; 4 for all timepoints for 10^5^ WT mice; and 3 for all timepoints for 10^3^ WT mice. **c**, Percentage of LS-DRG neurons infected by 6 dpi in tissue sections from 10^5^ PFU-inoculated hACE2 and WT mice. 42% of neurons were infected in hACE2 mice and 24% were infected in WT mice. **d**, Immunofluorescence for SARS-N (grey) and NeuN (red) in LS-DRG sections from infected and uninfected hACE2 and WT mice at 6 dpi. SARS-N is more prevalent in hACE2 than in WT but observable in both. No SARS-N was detected in uninfected mice. Detection of RNA and SARS-N in peripheral neurons with no direct connection to the oronasopharynx suggests spread via hematogenous dissemination or via axonal transport. **e**, Immunofluorescence for SARS-S (grey) and NeuN (red) in LS-DRG sections from infected hACE2 and WT mice at 6 dpi. SARS-S immunostaining was similar to that of SARS-N, with neuronal staining in hACE2 mice and satellite glial cell (SGC) staining in WT mice. **f**, Immunofluorescence for dsRNA (grey) and NeuN (red) in LS-DRG sections from infected hACE2 and WT mice at 6 dpi. dsRNA immunostaining was similar to that of SARS-N and SARS-S with neuronal staining in hACE2 mice and SGC staining in WT mice. Presence of dsRNA indicates viral genome replication. **g**, SARS-N (grey) was detected in numerous satellite glial cells (SGCs, glutamine synthetase (GS), red) surrounding infected LS-DRG neurons. See Supplementary Video 1 for 3D rendering of this image. **h**, Representative image of immunofluorescence for SARS-N and NeuN in spinal cord cross-sections from an infected hACE2 mouse at 6 dpi. SARS-N was observed as discrete puncta (arrowheads in e1 and e2) in neuronal cytoplasm, reminiscent of viral replication complexes. See Supplementary Video 2 for 3D rendering of this image **i**, Representative image of immunofluorescence for SARS-N (grey) and microglial marker Iba1 (red), in spinal cord cross-sections from an infected hACE2 mouse at 6 dpi. Microglia processes are present throughout the cord as are discrete SARS-N puncta. **j**, Representative image of immunofluorescence for SARS-N (grey) and astrocyte marker S100B (red), in spinal cord cross-sections from an infected hACE2 mouse at 6 dpi. Astrocytes are present throughout the cord as are discrete SARS-N puncta. Detection of RNA and SARS-N in the spinal cord demonstrates central neurons are susceptible to infection and may be infected directly from the LS-DRG, brainstem, or hematogenously. Data are the mean ± s.e.m. Log-transformed RNA genome copy numbers were statistically compared by three-way ANOVA (independent variables: inocula, days post infection, genotype). Pairwise comparisons were conducted using Tukey’s HSD post hoc tests. \**p* < 0.05. See Extended Data Figure 2 for unmerged LS-DRGs. See Extended Data Figure 3 for unmerged spinal cord sections and controls. See Supplementary Figure 1 for additional antibody validation via western blot, hACE2 genotyping, hACE2 protein expression, and RT-qPCR controls.

### Individual brain regions support varying levels of viral invasion and reproduction

Previous studies have tested brain homogenates to assess viral RNA and infectious virus in the brain, which does not allow for spatial analysis of and may obscure detection of low levels of virus in specific brain regions^14, 16, 21–23^. To determine if SARS-CoV-2 is present in specific brain regions, we assessed the olfactory bulb, hippocampus, cortex, brainstem, and cerebellum. We detected viral RNA in all brain regions at 3 dpi, which increased by 6 dpi, in both hACE2 and WT mice (Fig. 3a,b,c,d,e). Higher viral RNA concentrations were found in hACE2 mice compared to WT mice in all regions, with the highest in hippocampus and brainstem at 6 dpi. In WT mice, all regions contained similar quantities of SARS-CoV-2 RNA, suggesting that ACE2-independent spread or replication through the nervous system may differ from hACE2 mice. Infectious virus was recovered from the hippocampus and brainstem (3 log PFU/mg homogenate) as early as 3 dpi in hACE2 mice and from all regions by 6 dpi, with highest concentrations in hippocampus and cortex (5 log PFU/mg homogenate) followed by olfactory bulb and brainstem (4 log PFU/mg homogenate) and cerebellum (2 log PFU/mg homogenate) (Fig. 3f). Unexpectedly, infectious virus was recovered from the hippocampus (10^3^ group: 2 PFU/mg homogenate; 10^5^ group: 5 PFU/mg homogenate) and brainstem (10^5^ group: 3 PFU/mg homogenate) from WT mice at 6 dpi. These results indicate that viral invasion and replication in the brain is not uniform, and that the highest concentrations of virus are found in areas connected to the olfactory and limbic systems, which are strongly affected during COVID-19. To further assess localization of SARS-CoV-2 in brain regions, we immunostained sagittal sections of hACE2 and WT brains for SARS-N. In the hippocampus (Fig. 3g), neurons with cytoplasmic and axonal SARS-N were evident in the majority of CA1 and CA3 neurons, which receive input from the olfactory system through the entorhinal cortex, supporting axonal spread to the hippocampus. At 6 dpi, the majority of neurons in the frontal cortex, lateral preoptic area and visual cortex, thalamus and nucleus accumbens were SARS-N-positive (Fig. 4, Extended Data Fig. 3). Some of these regions also showed diffuse SARS-N in tissue (cortex, reticular formation), suggesting differences in pathology within various brain regions. As the reticular formation facilitates motor activity associated with the vagus nerve, pathology in this region would correlate with vagus nerve dysfunction, which is associated with even mild COVID-19 cases^24^. Although few SARS-N-positive cells were found in the olfactory bulb, abundant cytoplasmic and axonal staining in the olfactory tubercle suggests axonal spread from the olfactory bulb into the tubercle. While immunostaining of WT brains was unremarkable (Extended Data Fig. 4), as previously published, viral RNA and infectious virus was readily recoverable, thus demonstrating the utility of using multiple complementary assays to assess the presence of neuroinvasive virus.

**Fig 3.**
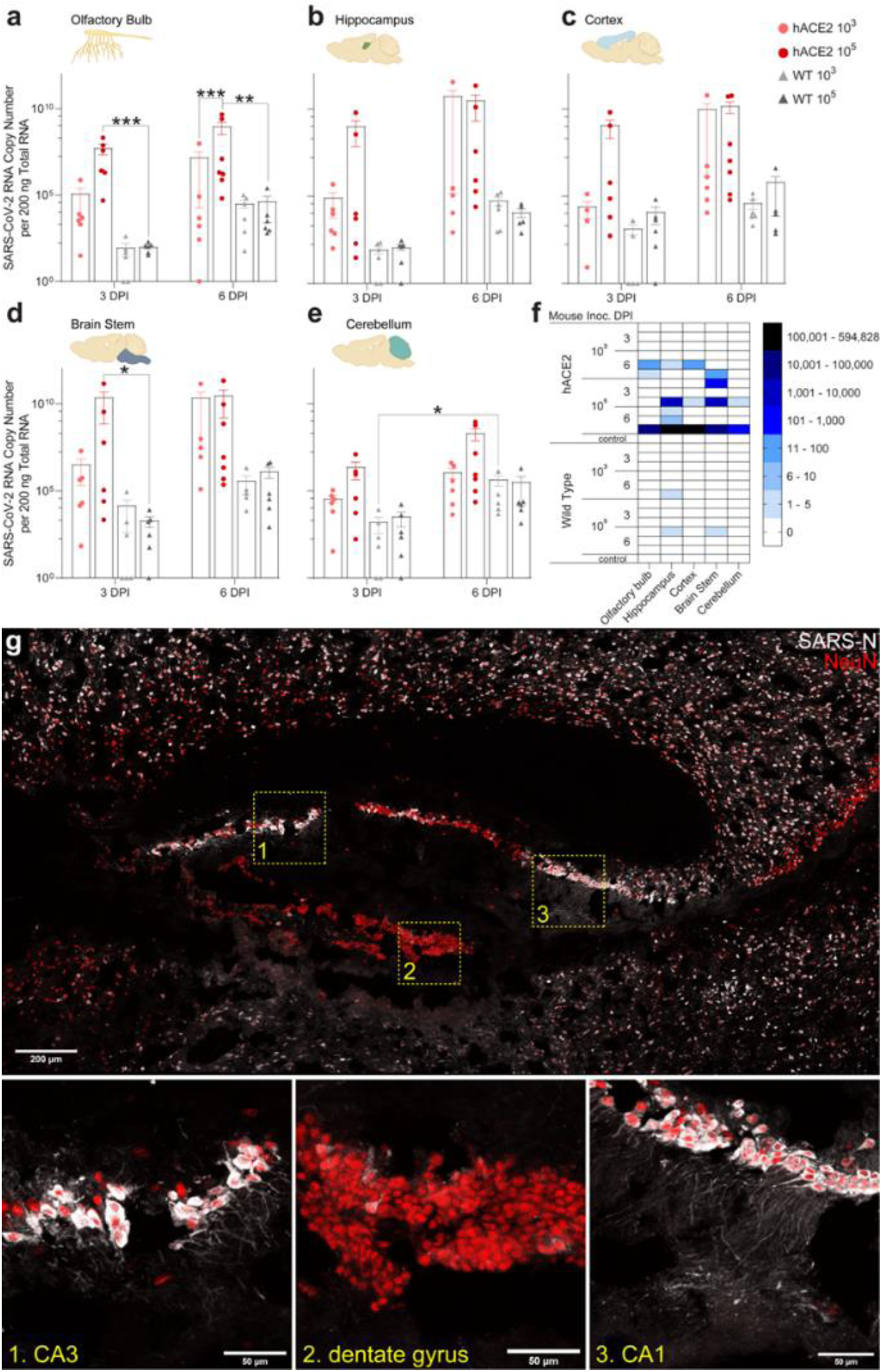
SARS-CoV-2 infection of the olfactory bulb and various brain regions in hACE2 and WT mice. SARS-CoV-2 RNA was detected in increasing concentration from 3-6d pi in olfactory bulb **a**, hippocampus **b**, cortex **c**, brainstem **d**, and cerebellum **e**, of hACE2 and WT mice in both inoculum groups. Three-way ANOVA detected a significant difference (F(7, 41) = 11.825, P = <0.0001) in RNA genome copy number in the olfactory bulb. Tukey’s HSD detected differences in the olfactory bulb between hACE2 and WT groups inoculated with 10^5^ PFU assessed at 3 dpi (*p*<0.0001) as well as between those groups assessed at 6 dpi (*p=*0.004). A significant difference (F(7, 41) = 5.433, P = 0.0002) was also detected in the hippocampi by three-way ANOVA, however Tukey’s HSD revealed it occurred between non-biologically relevant comparisons. A significant difference (F(7, 41) = 7.217, P = <0.0001) was also detected in the brainstem of the hACE2 and WT groups inoculated with 10^5^ PFU assessed at 3 dpi (*p*= 0.0107). While differences were detected in the cortex (F(7, 41) = 6.302, P = <0.0001) none were between relevant groups. A significant difference was detected in the cerebellum (F(7, 41) = 6.996, P = <0.0001) of the WT groups inoculated with 10^3^ PFU when compared between 3 and 6 dpi. **f**, Heatmap showing recovery of infectious SARS-CoV-2 from homogenates of the olfactory bulb and specific brain regions assessed for viral RNA. Recovery of infectious virus varied across individual animals with some having no regions with recoverable virus and some with virus in all regions. Of note, infectious virus was recovered from the hippocampi (5 PFU/mg homogenate) and brainstem (3 PFU/mg homogenate) of some WT mice, which are regions impacted in COVID-19 disease. **g**, Immunofluorescence for SARS-N (grey) and NeuN (red) in the hippocampus from an infected hACE2 mouse assessed 6 dpi. SARS-N+ neurons were observed in CA3 and CA1. Very few SARS-N+ neurons were detected in the granule cell layer of the dentate gyrus. Of note are SARS-N+ processes in infected neurons throughout the hippocampus, as shown in the magnified regions in lower panels. Data are the mean ± s.e.m. Log transformed RNA genome copy numbers were statistically compared by three-way ANOVA (independent variables: inocula, days post infection, genotype). Pairwise comparisons were conducted using Tukey’s HSD post hoc tests. \**p* < 0.05, \*\**p* < 0.01, \*\*\**p* < 0.001. n=6 for all animals/timepoints/tissues (except 10^5^ hACE2 6 dpi brain regions: n=7). See Extended Data Figure 3 for control hACE2 brain sections and Extended Data Figure 4 for infected and control WT brain sections. See Supplementary Figure 1 for additional antibody validation via western blot, hACE2 genotyping, hACE2 protein expression, and RT-qPCR controls.

**Fig 4.**
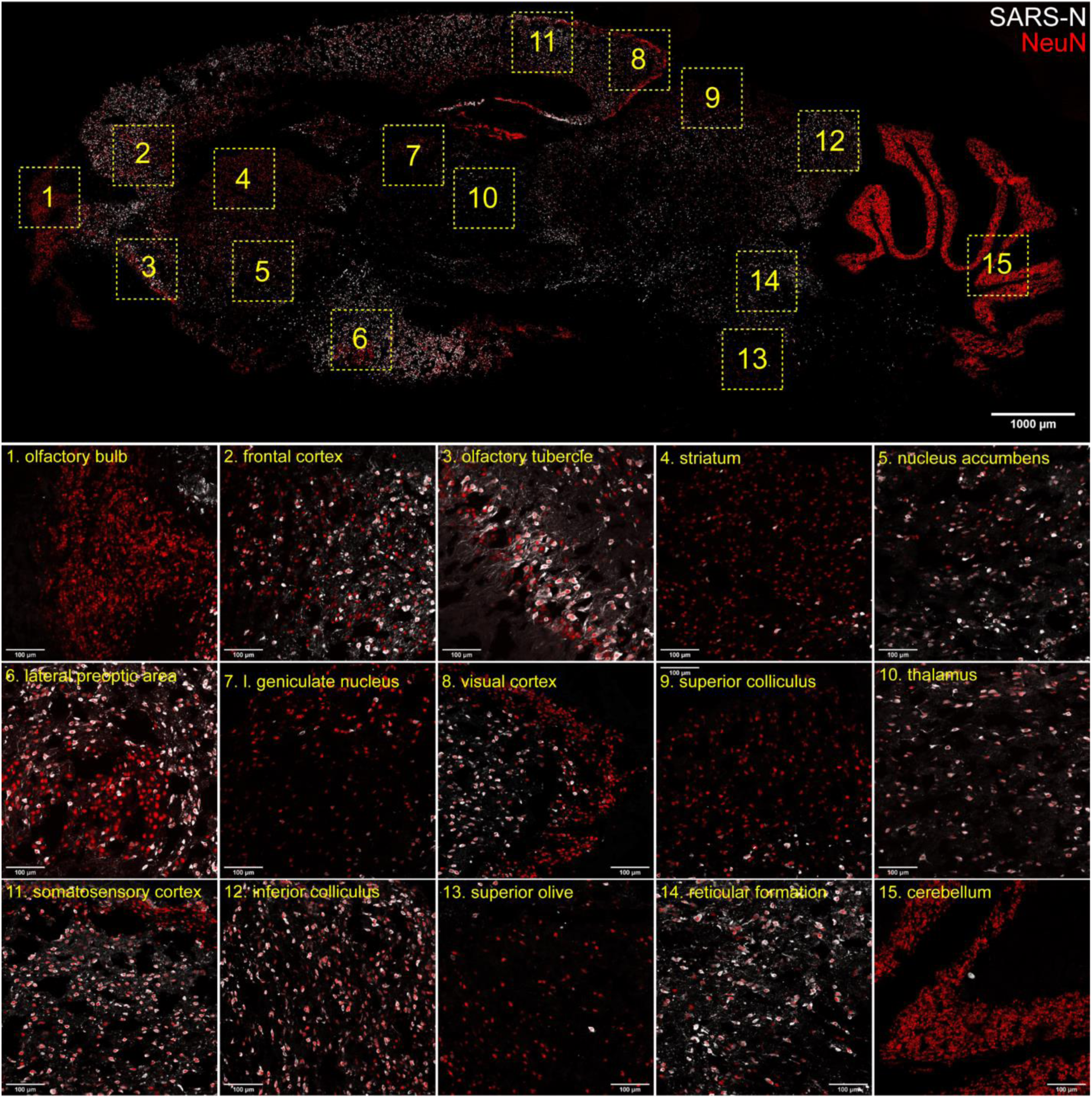
Immunofluorescence for SARS-N (grey) and NeuN (red) in a parasagittal brain section from an infected hACE2 mouse assessed 6 DPI. SARS-N+ neurons were observed throughout multiple brain regions with some notable exceptions. Few SARS-N+ cells were detected in the granule cell layer of the olfactory bulb, the medium spiny neurons in the striatum, and the granule cells in the cerebellum. Relatively few SARS-N+ neurons were identified in the lateral geniculate nucleus, layers 2 and 3 of the visual cortex, and in the superior colliculus. This stood in contrast to adjacent non-visual areas (thalamus, somatosensory cortex, inferior colliculus), which had numerous SARS-N+ neurons. The fact that some of these areas contain GABAergic neurons suggests that inhibitory neurons may be spared infection, at least at this time point. See Extended Data Figure 3 for control hACE2 brain sections and Extended Data Figure 4 for infected and control WT brain sections. See Supplementary Figure 1 for additional antibody validation via western blot, hACE2 genotyping, hACE2 protein expression, and RT-qPCR controls.

### SARS-CoV-2 productively infects primary cultured sensory but not autonomic neurons from adult mice

To confirm that PNS sensory and autonomic neurons are both susceptible and permissive to infection with SARS-CoV-2, resulting in release of infectious virus, and to establish basic replication kinetics of SARS-CoV-2 in neurons, primary neuronal cultures were established from LS-DRGs, TGs, and SCGs from 8-10-week-old hACE2 and WT mice. Following infection, media and cells were analyzed separately for viral RNA and infectious virus to differentiate between intracellular replication and release of infectious virus. Viral RNA levels increased in SCG neurons of hACE2 and WT mice, although the increase was mostly in cells, not media (Fig. 5a). Infectious virus was not detected in cellular or media fractions (Fig. 5b). While viral genome replication occurred in SCGs, infectious virus was not released, suggesting abortive infection in sympathetic autonomic neurons. Considering the pathology of the SCGs *in vivo*, SARS-CoV-2 appears to be cytotoxic to SCG neurons prior to production of viral progeny. In sensory neurons, viral RNA increased in TG and LS-DRG neurons in cyclical 48hr intervals, in both cellular and media fractions, with the exception of WT LS-DRG neurons, in which RNA steadily increased ultimately reaching concentrations similar to hACE2 neurons (Fig. 5a). Similarly, infectious virus was recovered cyclically at 48hr intervals from TG and LS-DRG neurons and media from hACE2 mice, but only a single cycle at 2 dpi in WT neurons (Fig. 5b). These data indicate successive rounds of genome amplification and infectious virus release occur in sensory neurons of the TG and LS-DRG, either within individual neurons without killing them or in additional neurons through a second infection cycle, although sustained production of viral progeny is dependent on hACE2.

**Fig 5.**
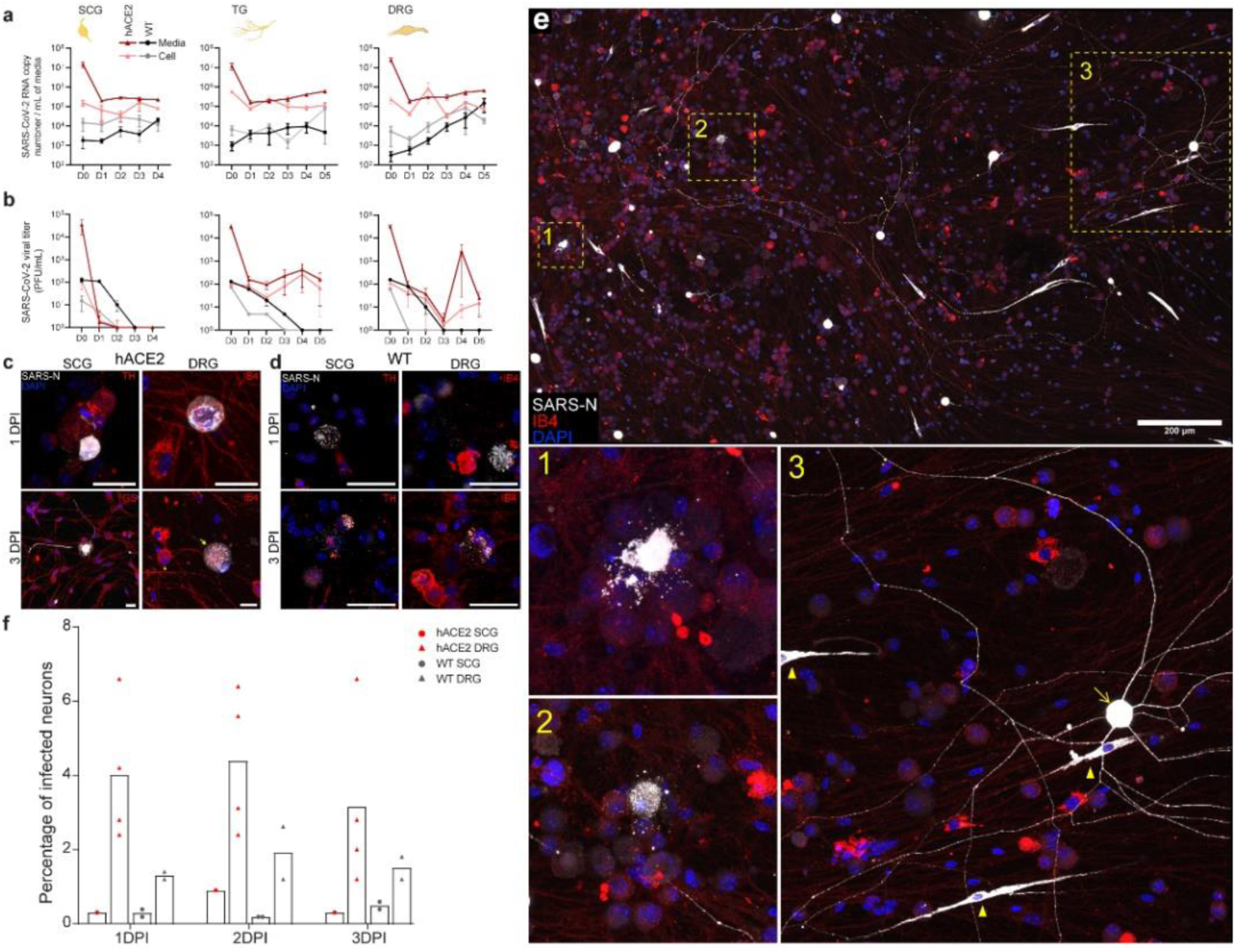
SARS-CoV-2 infection of primary neuronal cultures from SCG, TG, LS-DRG of hACE2 and WT mice. Primary neuronal cultures were generated from SCG, TG, and LS-DRG from 8-10 wk old hACE2 and WT mice. **a**, SARS-CoV-2 RNA was quantified by RT-qPCR separately in neuronal cells and media to generate a 4–5-day viral genome replication profile. Intracellular replication patterns are similar between hACE2 and WT neurons, although at reduced levels in WT neurons, with increasing viral RNA detected in media over the course of infection. hACE2 LS-DRGs have peaks in genome replication ∼48 hpi and ∼96 hpi indicating successive rounds of replication. Results are from three separate neuronal cultures, each with duplicate technical replicates per ganglion/timepoint. **b**, Infectious virus was quantified by plaque assay on Vero E6 cells from cellular and media fractions of SCG, TG, and LS-DRG neuronal cultures to generate growth curves in primary neurons from hACE2 and WT mice. Infectious virus was not recovered from SCG neurons indicating abortive infection, likely mediated by cytotoxicity. Infectious virus was recovered from TG and LS-DRG neurons indicating productive infection of these neurons, although sustained production of viral progeny is dependent on hACE2. Results are based on sample sizes as reported for above for panel a. **c**, hACE2 neuronal cultures: immunofluorescence for SARS-N (grey) and either tyrosine hydroxylase (TH, red) or Isolectin-B4 (IB4, red) to counterstain SCG and LS-DRG neurons, respectively, or glutamine synthetase (GS, red) to stain satellite glial cells. SARS-N was observed in neurons from each of the ganglia. Infected neurons were largely free of neurites by 1 dpi. At 3 dpi, many infected neurons exhibited cytopathologies such as degraded neurites, enlarged multi-nucleated cell bodies (arrow) compared to uninfected neurons (arrowhead), and SARS-N+ puncta reminiscent of viral replication compartments. See Supplementary Video 3 for 3D rendering of LS-DRG at 3 dpi. See Supplementary Video 4 for 3D rendering of TG at 2 dpi. **d**, WT neuronal cultures: immunofluorescence for SARS-N (grey) and either TH or IB4 (red) to counterstain neurons, or GS to stain satellite glial cells. Immunostaining revealed a similarly heterogenous infection of neurons and satellite glial cells as observed in hACE2 neurons. **e**, Immunofluorescence for SARS-N (grey) and IB4 to counterstain hACE2 LS-DRG neurons at 3 dpi shows a variety of phenotypes of infected cells, including neurons with a loss of membrane integrity (inset 1), SARS-N+ puncta within and surrounding neurons (inset 2), and seemingly healthy neurons with extensive neurites with strong SARS-N+ staining (arrow in inset 3). Infected satellite glial cells were also observed (arrowheads in inset 3); many appeared to be activated, noted by the presence of extended cellular processes. These findings are similar to immunostaining of LS-DRGs of hACE2 and WT mice *in vivo*, which also contained numerous infected satellite glial cells. **f,** Percentage of hACE2 autonomic (SCG) and sensory (LS-DRG) cultured neurons positive for SARS-N were counted from 1-3 dpi. A small percentage of autonomic (SCG) neurons were visibly infected, with significant observable cell death, similar to *in vivo* observations. Infection in sensory (LS-DRG) neurons were consistent from 1-3 dpi, with ∼5% infected. Infection of neurons *ex vivo* is less efficient than infection *in vivo*. Scale bar = 20 μm. Data are the mean ± s.e.m. See Extended Data Figures 5-6 for unmerged and control images. See Supplementary Figure 1 for additional antibody validation via western blot, hACE2 genotyping, hACE2 protein expression, and RT-qPCR controls.

In parallel, primary neuronal cultures were infected and fixed 1, 2, and 3-dpi for SARS-N immunostaining. In addition to genome replication, SCG neurons were permissive for SARS-N protein expression, shown by positive immunofluorescence in both hACE2 (Fig. 5c, Extended Data Fig 5) and WT neurons (Fig. 5d, Extended Data Fig 6), with activated SGCs accompanying dying neurons by 3 dpi (Fig. 5c,d). In LS-DRG neurons, several phenotypes were observed in both hACE2 (Fig. 5c, Extended Data Fig 5) and WT neurons (Fig. 5d, Extended Data Fig 6), including perinuclear SARS-N staining (1 dpi) and punctate staining in the cytoplasm of enlarged neurons (3 dpi), likely representing replication compartments (Supplementary Video 3-4). Infected LS-DRGs (Fig. 5e) showed a variety of phenotypes, including loss of membrane integrity (inset 1), cytoplasmic puncta (inset 2), and seemingly healthy neurons strongly expressing SARS-N in cytoplasm and processes (inset 3). Infected SGCs, which appeared to be activated, were also present in primary neuronal cultures (Fig. 5e inset 3, yellow arrowheads). The modest increase in viral RNA and infectious virus in media (Fig. 5a,b) can be explained by the small percentage of LS-DRG neurons (∼5% hACE2, ∼2% WT) and SCG neurons (<1% hACE2 or WT) that became productively infected in culture (Fig. 5f). Taken together, these data show that PNS sensory neurons are both susceptible and permissive to SARS-CoV-2 infection, resulting in release of infectious virus. Also, while autonomic neurons are susceptible to infection with SARS-CoV-2 and genome replication can occur, they are not permissive to release of infectious virus.

### Neuroinvasion of the PNS and CNS occurs before viremia

To determine if neuroinvasion is driven by hematogenous entry or direct neuronal entry, PNS and CNS tissues were assessed 18 and 42 hpi after intranasal inoculation of hACE2 and WT mice (Fig. 6a,b). Although no viral RNA was detected in blood at 18 hpi, viral RNA was detected in all PNS, the majority of CNS, and salivary gland (innervated by SCG) tissues in both hACE2 and WT mice. By 42 hpi, viremia was detected in a single hACE2 mouse, and viral RNA had increased in all hACE2 PNS and CNS tissues except SCGs. In WT mice, viremia was not detected and viral RNA was no longer detected in salivary glands, SCGs, or LS-DRGs, but had increased in brainstem and hippocampus by 42 hpi. Immunostaining did not detect SARS-N in any tissues, indicating the virus was transiting through PNS tissues when collected but had not yet begun replication.

**Fig 6.**
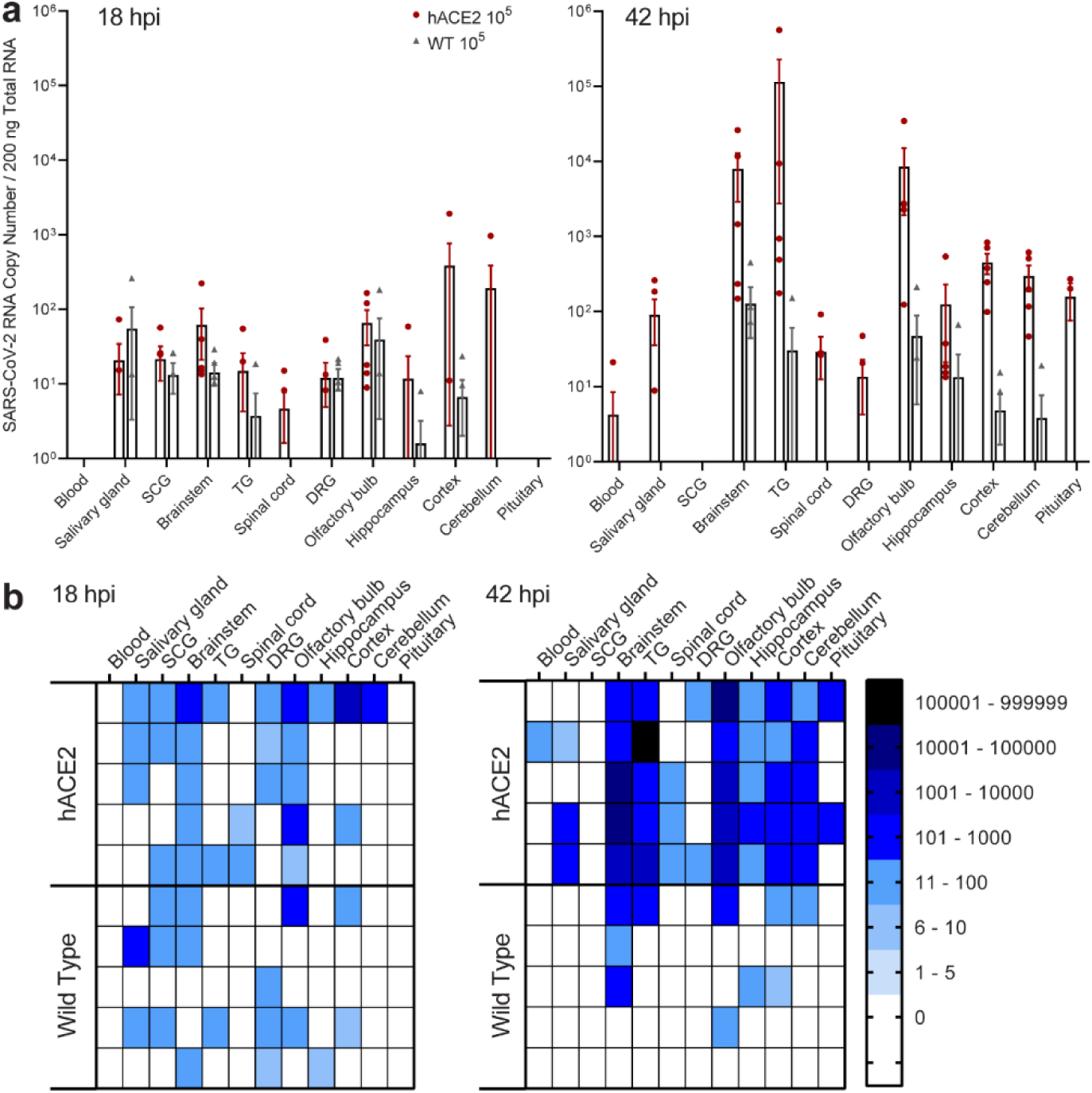
Pre-viremic neuroinvasion of SARS-CoV-2 into PNS and CNS of hACE2 and WT mice as early as 18 hours post infection. **a,** Although no viral RNA was detected in blood, low levels of SARS-CoV-2 RNA were detected in PNS and CNS of both hACE2 and WT mice as early as 18 hpi. CNS invasion in hACE2 and WT mice was common in olfactory bulb, hippocampus, cortex, and brainstem. Viral RNA was detected in PNS ganglia and the tissues they innervate (SCG-salivary gland, TG-brainstem) in both hACE2 and WT mice. Viral RNA was detected separately in the LS-DRGs and spinal cords of some mice. By 42 hpi, viral RNA was detected in blood in only one hACE2 mouse, but had increased in brainstem, TG and olfactory bulb, indicating replication in these tissues. Sample sizes were hACE2 mice n=10 (5 per timepoint) and WT mice n=10 (5 per timepoint). **b**, Heatmaps visually displaying RT-qPCR values from panel a. Neuroinvasion in both PNS and CNS occurs rapidly before detectable viremia, thereby indicating direct neural entry and trans-synaptic spread of SARS-CoV-2. Detection of viral RNA in the LS-DRGs but not the spinal cord in some mice and vice versa in others indicates separate entry routes exist for the LS-DRG and spinal cord. Invasion of the cord likely occurs from the brainstem, as all mice with early spinal cord infection also had brainstem infection. Invasion of the LS-DRG may occur from the periphery. Infectious virus was not detected by plaque assay, indicating virus had not yet started replicating at these early time points. See Supplementary Figure 1 for hACE2 genotyping, hACE2 protein expression, and RT-qPCR controls.

To verify that these results were reproducible in an alternative non-transgenic animal model, golden Syrian hamsters were infected (n=9) and tissues collected 18 hpi, 42 hpi, and 3 dpi (Fig. 7a,b,c, Extended Data Fig. 1 a,b). Consistent with mice, viral RNA was detected in all PNS and CNS tissues at 18 hpi, preceding viremia, with continual increase at 42 hpi, indicating direct neural invasion independent of viremia (Fig. 7a,b). Unlike in mice, immunostaining showed presence of SARS-N in TGs, LS-DRGs, and SCGs at 18 hpi, with continued presence through 3 dpi. Substantial vacuolization was observed in the SCGs at 3 dpi, similar to pathology observed in mice (Fig. 7c). SARS-N staining was detected in neurons of infected hamster brains (Extended Data Fig 7, Supplementary Video 5). In light of LS-DRG/spinal cord infection observed in mice, the functional impact of infection on the sensory nervous system was assessed. Using the von Frey assay, a significant decrease (*p*=0.0001) in the amount of pressure required to elicit a withdrawal reflex was noted, demonstrating allodynia (Fig. 7d). Allodynia occurred rapidly as a consequence of infection; 55% (n=5 of 9) of hamsters demonstrated allodynia by 18 hpi and all remaining hamsters (n=3 of 3) by 3 dpi. Thus, neuroinvasion occurs rapidly after infection, is mediated by invasion of and transport along neurons, can occur independent of hACE2, and functionally impacts sensory neurons resulting in allodynia. Other studies have shown transcriptional changes in the CNS and to a lesser extent the PNS (TG), as early as 3 dpi following SARS-CoV-2 infection of hamsters^25^. Our results show neuroinvasion via direct neuronal entry into the SCG, TG, and DRG within 18 hpi following infection, providing a mechanism for dysregulation of neuronal gene expression.

**Fig 7.**
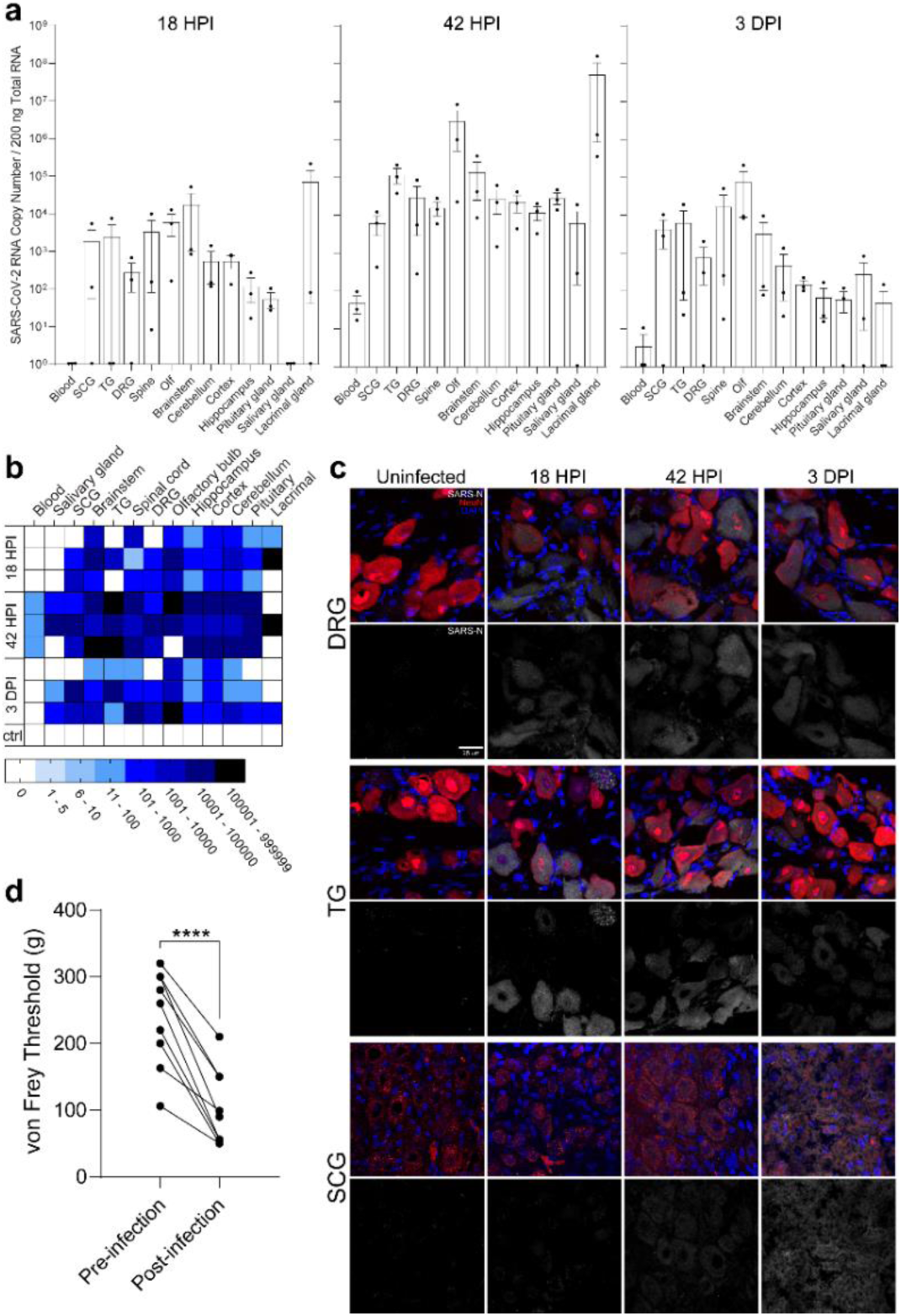
Pre-viremic neuroinvasion of SARS-CoV-2 into PNS and CNS in Syrian golden hamsters as early as 18 hours post infection. **a,** Similar to hACE2 and WT mice, no viral RNA was detected in hamster blood at 18 hpi, however low levels of SARS-CoV-2 RNA were detected in PNS and CNS tissues. Viral RNA was detected in PNS ganglia and the tissues they innervate (SCG-lacrimal gland, TG-brainstem, LS-DRG-spinal cord) as early as 18 hpi. By 42 hpi, viral RNA was increased in all PNS tissues (excluding SCGs), indicating replication in these tissues. Lacrimal glands harbored notably high viral RNA concentrations. Viral RNA copy number was stable in SCGs at 3 dpi (4 log), and had returned to levels observed at 18 hpi in TG and LS-DRGs (4 log and 3 log, respectively), indicating a burst of viral replication. Sample size was n=9 (3 hamsters per timepoint). **b**, Heatmaps visually displaying RT-qPCR values from panel a. Neuroinvasion in both PNS and CNS occurs rapidly before detectable viremia, thereby indicating direct neural entry and trans-synaptic spread of SARS-CoV-2, as observed in mice. **c**, Immunofluorescence for SARS-N (grey) and NeuN (red) in LS-DRG, TG, and SCG sections from infected and uninfected hamsters at 18 hpi, 42 hpi, and 3 dpi. SARS-N appears in all ganglia by 18 hpi and continues to be present to 3 dpi. Significant pathology/vacuolization is observed in SCGs at 3 dpi, similar to the pathology observed in mice. **d**, Results of von Frey threshold test showing a decrease in the amount of force required to elicit a withdrawal reflex, indicating allodynia as a consequence of infection. A paired samples two-tailed t-test found this decrease to be significant (t(8)=7.606, *p=*0.0001). Presence of SARS-CoV-2 neuroinvasion of LS-DRGs likely mediates this allodynia, which was noted in some animals (5 of 9) as early as 18 hpi and all remaining animals (3 of 3) by 3 dpi. Scale bar = 20 μm. Data are the mean ± s.e.m. See Extended Data Figure 7 for immunostaining of infected hamster brain. See Supplementary Figure 1 for additional antibody validation via western blot and RT-qPCR controls.

### Neuronal entry involves neuropilin-1 (NRP-1)

NRP-1 has been shown to interact with SARS-S, thereby enhancing viral binding/entry in non-neuronal cells^26–28^. Since our WT mice and hamsters were infected despite absence of hACE2, we investigated the contribution of NRP-1 to entry in primary sensory neurons. NRP-1 expression was confirmed in SCG, TG, and LS-DRG neurons of hACE2 and WT mice by western blot (Fig. 8a, Supplementary Fig. 1d,e). Immunostaining in LS-DRG neurons showed NRP-1 at cell membranes of neurons and SGCs (Fig. 8b). Primary cultured LS-DRG neurons from hACE2 and WT mice were pretreated with EG00229, a selective NRP-1 antagonist, infected with SARS-CoV-2, and viral RNA concentrations were assessed 2 dpi. Viral RNA concentrations were significantly reduced by 99.8% in hACE2 neurons (*p*=0.0081) and 86.7% in WT neurons (*p*=0. 0141; Fig. 8c), indicating that NRP-1 is a SARS-CoV-2 co-receptor in neurons irrespective of hACE2 expression. NRP-1 inhibition was repeated using escalating concentrations of a polyclonal NRP-1 blocking antibody (1, 15, 30 µg). Viral RNA concentrations were significantly reduced by 99.62% (*p*=0.0123) in hACE2 neurons treated with 30 µg antibody (Fig. 8d). Interestingly, at this concentration the blocking antibody enhanced entry in WT neurons.

**Fig 8.**
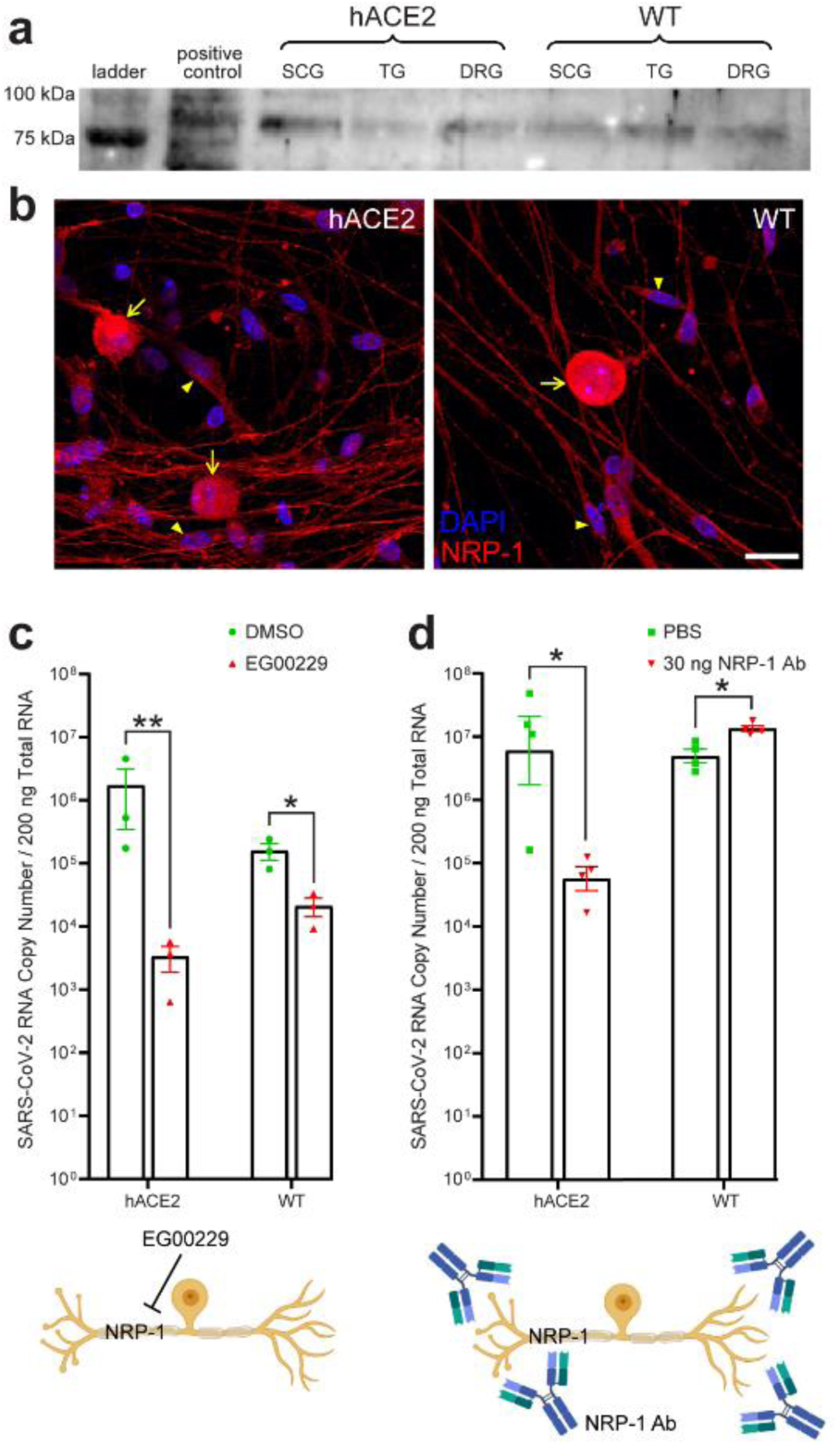
The role of NRP-1 during neuronal entry of SARS-CoV-2. **a,** Western blot of ganglia (SCG, TG, LS-DRG) from hACE2 and WT mice showing the presence of the transmembrane glycoprotein neuropilin-1 (NRP-1) in ganglia homogenates. NRP-1 has been shown to increase SARS-CoV-2 entry into non-neuronal cells. **b**, Immunofluorescence for NRP-1 (red) and DAPI (blue) in primary neuronal cultures of LS-DRG neurons from uninfected hACE2 and WT mice, showing that NRP-1 is present across the soma and processes of neurons as well as satellite glial cells. **c**, Treatment of hACE2 and WT LS-DRG neurons with the NRP-1 antagonist EG00229 prior to infection with SARS-CoV-2 significantly reduced viral RNA concentrations at 2 dpi, the initial peak of viral replication in hACE2 LS-DRG neurons as determined through our replication/growth curves, by 99.8% (t(4) =4.896, *p*=0.0081) in hACE2 neurons and 86.7% (t(4) =4.165, *p*=0.0141) in WT neurons. **d**, Treatment of hACE2 and WT LS-DRG neurons with a polyclonal NRP-1 blocking antibody (AF566, R&D Systems) prior to infection with SARS-CoV-2 significantly reduced viral RNA concentrations at 2 dpi by 99.6% in hACE2 LS-DRG neurons (t(6) =3.535, *p*=0.123), but increased viral RNA concentrations by 53.2% in WT LS-DRG neurons (t(6) =3.701, *p*=0.101). The difference in results between EG0029 and the NRP-1 blocking antibody is likely mediated by the site of action of each. EG00229 putatively blocks binding between the carboxyl-terminal sequence of SARS-CoV-2 S1 which has a C-end rule (CendR) motif and the extracellular b1b2 CendR binding pocket of NRP-1, which has been suggested as an alternative co-receptor for SARS-CoV-2 in non-neuronal cells. The mechanism of action of EG00229 is therefore likely more specific in its blockade of the NRP-1/SARS-CoV-2 spike protein binding site than that of a polyclonal NRP-1 blocking antibody. Thus, NRP-1 is a critical co-receptor mediating viral entry into neurons expressing hACE2 and also enhances viral entry into WT neurons. These data also indicate that that additional host proteins are involved in neuronal entry. Scale bar = 20 μm. Data are the mean ± s.e.m. Results are from two separate neuronal cultures, each with duplicate technical replicates per ganglion/NRP-1 inhibitor type. See Supplementary Figure 1 for additional antibody validation via western blot, hACE2 genotyping, hACE2 protein expression, NRP-1 protein expression, and RT-qPCR controls.

## Discussion

Neurotropic viruses can enter the nervous system hematogenously or through neural pathways. They can infect vascular endothelial cells to access underlying tissues, or transport across inside extravasating leukocytes. Viruses can enter neural pathways through peripheral sensory, autonomic, and/or motor axon terminals and reach the CNS by retrograde transport, often moving trans-synaptically along functionally connected pathways. SARS-CoV-2 likely uses both mechanisms. SARS-CoV-2 CNS invasion has been proposed via infection of OSNs in the nasal neuroepithelium, the olfactory bulb, and its cortical projections^14–16, 19, 21–23^. Organoid, stem cell, microfluidic, and mouse models, correlating with human autopsy findings, demonstrate disruption of endothelial barriers and choroid plexus integrity, as well as transcytosis of SARS-CoV-2, supporting hematogenous CNS entry^5, 19, 29–33^. While OSNs are a key constituent of the nasal neuroepithelium, the oronasopharynx is innervated by other sensory and autonomic pathways through which SARS-CoV-2 may enter the nervous system. Utilizing hACE2 mice, WT mice, golden Syrian hamsters, and primary neuronal cultures, we show susceptibility of peripheral neurons to SARS-CoV-2 infection, demonstrating differential replication kinetics and cytopathic outcomes following infection of sensory, autonomic, and central neurons. We also show evidence supporting axonal transport of SARS-CoV-2 and CNS entry, preceding viremia, through neural pathways that functionally connect to brain regions responsible for memory and cognition. Furthermore, we show that SARS-CoV-2 can use NRP-1 for neuronal entry in the absence of hACE2. As COVID-19 neurological symptoms are often related to peripheral neuron dysfunction, focusing solely on CNS neuroinvasion overlooks the potential impacts of SARS-CoV-2 in the PNS.

The detection of SARS-CoV-2 in TGs and SCGs is consistent with their innervation of the oronasal mucosa. Both animal models showed similar viral RNA concentrations 18 hpi in the TG, SCG, and olfactory bulb, indicating sensory and autonomic pathways are as susceptible to invasion as the olfactory system. Our results complement findings from post-mortem studies of COVID-19 patients, which have found viral RNA in the TGs of 14% of patients assessed^32^, with axonal damage and neuron loss in the TGs of some patients^33, 34^. Our use of *in vivo* and *ex vivo* infections demonstrates that sensory trigeminal neurons are both susceptible and permissive to productive infection with SARS-CoV-2, culminating in release of infectious virus, suggesting the TG can serve as an alternative route for CNS invasion. Increasing reports of acute necrotizing hemorrhagic encephalopathy (AHNE) associated with COVID-19 suggest that neuroinvasion may follow a similar pathway as another neurotropic virus, HSV-1, which causes a nearly identical complication upon reaching the CNS from TGs. Invasion of the brainstem along projections of the TG could damage nuclei important in cardiorespiratory regulation, a feature of severe COVID-19, and at least one imaging study of a COVID-19 patient with AHNE suggested invasion of the brainstem via the TG^7^. While viral RNA and protein have been detected in salivary glands of COVID-19 patients, and infectious virus has been recovered from saliva, no assessments of the SCG have been made in human autopsies^35–38^. The SCG provides sympathetic nervous system innervation to the oronasal mucosa, salivary and lacrimal glands, and oronasopharynx vasculature, providing possible neural entry sites. Retrograde axonal transport along the SCG would allow for invasion of preganglionic neurons in the cervical spinal cord, shown to harbor SARS-CoV-2 RNA and protein at autopsy in some COVID-19 patients^39^. Our neuronal culture infection data indicate abortive infection in the SCG, suggesting that it may be of less concern for direct CNS invasion. However, the pathology of the SCG following infection *in vivo* in both animal models suggests that SARS-CoV-2 causes significant, and perhaps irreparable, damage to sympathetic neurons. Although we did not examine thoracic ganglia or the sympathetic trunk, the cytopathology in the SCG has broader implications for cardiac function, which relies on autonomic regulation. It is notable that comparable viral RNA concentrations were detected in TGs and SCGs of WT mice and hamsters, indicating that both tissues are equally susceptible to infection in the absence of hACE2.

Entry into TGs and SCGs following intranasal inoculation was expected, but detection of SARS-CoV-2 in LS-DRGs was unanticipated. Viral RNA in LS-DRGs at levels comparable to TGs in both animal models demonstrates that distal sensory neurons, regardless of their location, are equally susceptible to infection. How the virus reached such distal ganglia is uncertain. Given that viral RNA was detected in LS-DRGs 18 hpi, preceding viremia, hematogenous spread is unlikely. It is noteworthy that detection of viral RNA in the LS-DRG and spinal cord occurred asynchronously at 18 hpi. However, virus was detected in both LS-DRG and spinal cord by 3 dpi. All mice and hamsters with early spinal cord infection also had brainstem infection. Thus, entry into spinal cord appears to follow CNS invasion at the brainstem but infection of LS-DRG neurons is possible from either central or peripheral axon terminals. Once in the LS-DRG, results from our primary neuronal cultures indicate that viral replication and release of infectious virus occurs cyclically every 48 h. As the virus infected a small percentage of sensory neurons in culture, this pattern likely represents two distinct cycles of productive infection. *In vivo*, however, the majority of LS-DRG neurons were positive for SARS-CoV-2, suggesting that infection of sensory neurons within the host is more efficient than *ex vivo* infection of cultured neurons. Although COVID-19 symptoms can include tingling, numbness, and burning in fingers and toes, which are indicative of nociceptor damage or dysfunction, and post-mortem studies of COVID-19 patients have found viral RNA in the sciatic nerve^39^, LS-DRG neurons have not previously been assessed for their susceptibility to infection with SARS-CoV-2, either in animal models or in human autopsies. However, ACE2 is expressed by a subset of nociceptors in human DRGs, particularly in LS-DRGs^40^. Development of allodynia in hamsters demonstrates that SARS-CoV-2 infection of peripheral sensory neurons results in sensory dysfunction. Spinal cord involvement is becoming increasingly associated with COVID-19 and a recent review of spinal cord disorders in COVID-19 posited that direct invasion of the cord by SARS-CoV-2 could cause these pathologies^41^. Our results indicate that neurons in the spinal cord and in the LS-DRG are susceptible to SARS-CoV-2 infection, which results in allodynia, providing a rationale for a deeper investigation of the LS-DRG as a site of productive viral infection in COVID-19 and the possibility of directional axonal transport.

Our findings reveal a more detailed picture of SARS-CoV-2 invasion into the brain than previous reports. We detected viral RNA and infectious virus in the hippocampus, cortex, and brainstem before viremia. TG neurons, with axonal projections to both oronasal epithelium and brainstem, or SCG neurons, with synaptic connectivity to salivary glands and brainstem, could deliver virus directly to the CNS. Viral RNA and protein have been detected in the olfactory bulb, brainstem, cerebellum, and cortex of COVID-19 patients^6, 42, 43^. Our results indicate that viral penetration and replication within the brain is region-specific, suggesting the presence of factors (cell types, synaptic connections, vascularization level) that favor invasion and replication in some regions over others. Our results comport with findings from post-mortem studies of COVID-19 patients that show brain region-dependent variation in viral RNA, including subgenomic RNA detection^39^. Our detection of infectious virus in the hippocampi and brainstems of both animal models emphasizes the importance of these structures in COVID-19 pathology. Notably, SARS-CoV-2 was detected in CNS and PNS of hACE2 mice, WT mice, and hamsters, indicating that hACE2 expression is not a strict requirement for neuronal infection. We demonstrate that NRP-1, previously shown to mediate entry into non-neuronal cells, also facilitates entry into neurons. Inhibition of NRP-1 in cultured LS-DRG neurons from hACE2 mice reduced infection to a greater extent than in WT neurons, indicating that NRP-1 can serve as a co-receptor to enhance infection in the presence of hACE2 expression or an alternative receptor independent of hACE2. The difference in results between EG00229, specific to the NRP-1/SARS-CoV-2 spike protein binding site, and the NRP-1 polyclonal blocking antibody is likely mediated by the site of action of each or influenced by other entry mediators.

The impact of SARS-CoV-2 on the nervous system is only beginning to be understood, with sensory and autonomic disorders lasting well beyond initial infection, indicating the need for a comprehensive understanding of its impact on the entire nervous system, not just the brain. Existing reports have described PASC symptoms involving the TG (trigeminal neuralgia)^44, 45^, SCG (Horner syndrome)^46^, LS-DRG (radicular pain)^47^, and spinal cord (transverse myelitis)^48^. Our *in vitro* and *in vivo* data indicate these tissues are susceptible to infection with SARS-CoV-2, with variable cytopathic outcomes, and merit investigation. No study has yet assessed the permissiveness of TG or SCG neurons to productive infection by SARS-CoV-2 or investigated their role in viral invasion of the CNS, though some post-mortem studies have detected viral RNA after protracted disease^33, 34^. Additionally, no studies have determined if distal sensory ganglia such as the LS-DRG can support infection, let alone provide an avenue of infection to the spine. We found that PNS sensory and autonomic neurons, spinal cord neurons, and supporting cells are susceptible and, in most cases, permissive to productive infection with SARS-CoV-2 via direct neural invasion, independent of hACE2, using NRP-1 as a co-receptor. We show that neuroinvasion is not a rare event following SARS-CoV-2 infection and that it occurs well before the onset of symptomatic disease. The rapidity of neuroinvasion severely limits the availability of autopsy samples to investigate early neuroinvasion events, as most samples are collected from patients well into their disease process. The presence of infectious virus in these tissues preceding viremia shows that neuroinvasion occurs via peripheral neural pathways. Further research into these sites of neuroinvasion is necessary with contemporary SARS-CoV-2 isolates, as COVID-19 transitions from a pandemic to an endemic disease associated with long-term neurological sequelae.

## Methods

### Ethics statement

This study was approved by the Virginia Polytechnic Institute & State University Institutional Animal Care and Use Committee (Protocol # 20-228, approved 01 Feb 2021 and Protocol #20-184, approved 02 Aug 2022). This study was carried out according to the US Department of Agriculture’s Animal Welfare Act and the Public Health Service’s Policy on Humane Care and Use of Laboratory Animals.

### Cells and virus

SARS-CoV-2 isolate USA-WA1/2020 (NR-52281; BEI Resources) was passaged twice using Vero E6 cells (CRL-1586, ATCC) by the Bertke Lab to produce viral stock for mouse and primary neuronal culture infections. USA-WA1/2020 was recovered from an oropharyngeal swab taken from a 35-year-old male in Washington state in January 2020 who was diagnosed with COVID-19 after returning from visiting family in Wuhan, China. Viral stocks were titrated in duplicate using a standard plaque assay on Vero E6 cells with agarose overlay^49^. Viral stock was sequenced by the VT-Molecular Diagnostics Laboratory (Fralin Biomedical Research Institute). Viral sequence data was deposited in GenBank (Accession# OP934235.1). Viral stocks for hamster infections were produced and titrated in Vero E6 cells using an independent aliquot of SARS-CoV-2 isolate USA-WA1/2020 (NR-52281; BEI) by the Duggal lab. Vero E6 cells and HEK293 cells (CRL-1573, ATCC) were maintained following standard cell culture protocols.

### Mouse infections

Eight to ten-week-old male and female B6.Cg-Tg(K18-ACE2)2Prlmn/J mice (Stock # 034860; Jackson Laboratory; n=12, 2 groups of 6 mice), and their wild-type C57BL/6J counterparts (n=12, 2 groups of 6 mice) were inoculated intranasally with SARS-CoV-2 isolate USA-WA1/2020 (Extended Data Fig 1a). Inoculations were carried out under ketamine/xylazine anesthesia in the on campus ABSL-3 facility after a one-day acclimation period. Mice received 20 µL of either 10^3^ PFU (2 groups per mouse type) or 10^5^ PFU (2 groups per mouse type) of SARS-CoV-2 in 1X PBS. The inoculum was split between the nares for each mouse. Uninfected K18-hACE2 control mice (n=2) and C57BL/6J wild-type control mice (n=2) were housed in a separate on campus ABSL-1 facility. Aliquots of the inocula and viral stock were saved for back titration using plaque assay for infectious viral titer and RT-qPCR for RNA copy number. All mice were genotyped using tail snip following Jackson Laboratory protocol # 38170 V2 (Extended Data Fig 1b). Mice were assessed daily for signs of disease and changes in weight and temperature. Mice from each group (K18-hACE2, WT) and inoculum dose (10^5^ PFU, 10^3^ PFU) were euthanized 3 days post infection (dpi) (n=3) and at 6 dpi (n=3). Tissues collected included blood, CNS tissues (olfactory bulb, hippocampus, brainstem, cerebellum, cortex, spinal cord), PNS tissues (autonomic ganglia: superior cervical ganglia-SCG; sensory ganglia: lumbosacral dorsal root ganglia-LS-DRG, trigeminal ganglia-TG), viscera (lung). Half of the tissues were collected in TRI Reagent for RNA extraction and RT-qPCR and the other half collected in 10% formalin for immunostaining. Brains were split into hemispheres maintaining attachment with the olfactory bulb. One hemisphere was fixed in formalin for immunostaining and the other dissected out into individual brain regions with each placed in TRI Reagent for RT-qPCR. This experiment was repeated as above for reproducibility, with the addition of an extra mouse in the 10^5^ PFU hACE2 6 dpi group to account for loss due to death. Tissue collection from the second experiment was split between TRI Reagent for RT-qPCR as above or flash frozen on dry ice for plaque assay. Blood samples from the initial and replicate infection study were assessed via RT-qPCR to assess for disseminated infection at 3- and 6-dpi. Lungs from the initial and replicate infection studies were assessed to verify infection at 3- and 6-dpi.

To assess viral spread through nervous tissues at earlier timepoints during infection, and to determine the role that viremia plays verses direct neuronal invasion, K18-hACE2 (n=10) and WT mice (n=10) were infected with 10^5^ PFU SARS-CoV-2 as described above and were euthanized 18- and 42-hpi (n=5 of each group/day). Blood, PNS tissues, CNS tissues (with addition of pituitary gland), and salivary glands were collected as described above for RT-qPCR, plaque assays, and immunostaining.

### Hamster infections and von Frey assessment of acute allodynia

To verify that results from the early timepoint neural invasion studies in mice were reproducible in an alternative animal model, and to assess the functional impact of infection on somatosensation, golden Syrian hamsters were infected using an independently propagated viral stock. After a two-day acclimation period in the ABSL-3, eight-week-old male golden Syrian hamsters (Stock #089; Envigo) were infected intranasally with 10^5^ PFU of SARS-CoV-2 isolate USA-WA1/2020 (n=9) under isoflurane anesthesia as previously described (Extended Data Fig 1a.)^50^. An uninfected control hamster (n=1) was housed in the ABSL-3 facility. Hamsters were monitored daily as described for mice. To assess for allodynia following infection of sensory neurons (LS-DRGs), which mediate transmission of sensory information from the periphery to the CNS, hamsters were tested daily using von Frey filaments on the hind paw footpad. Sensory testing began one day prior to infection, occurred on the day of infection preceding inoculation, and then occurred daily until euthanasia. A series of increasing caliber von Frey filaments were used on the footpad until a withdrawal reflex was elicited. von Frey filaments were cleaned between each animal. Post infection measurements were averaged for each animal and compared to the average of the pre infection measurements of the same animal by analysis of area under the curve. Hamsters were euthanized 18 hpi, 42 hpi, and 3 dpi (n=3 each day) as described above for mice. Tissue types collected, methods of collection, and downstream assays were the same as described for the early timepoint neural invasion studies in mice, with the addition of lacrimal glands.

### RNA extraction and SARS-CoV-2 specific RT-qPCR

RNA was extracted and RT-qPCR performed as previously described^51^. Briefly, tissues were homogenized in 200 µL TRI Reagent (Fisher Scientific) using a handheld tissue homogenizer with sterile pestles (Cole-Parmer). RNA was extracted using a standard guanidinium thiocyanate-phenol-chloroform extraction. RNA purity and quantity were assessed using a NanoDrop 2000 spectrophotometer (ThermoFisher). SARS-CoV-2 RT-qPCR reactions (10 µL) using the iTaq Universal Probe One-Step Kit (BioRad) and SARS-CoV-2 N1 primers/probe mix (Stock# 10006713; Integrated DNA Technologies) were run on a ViiA 7 Real-Time PCR system (Applied Biosystems) as described in the instructions for use of the CDC 2019-Novel Coronavirus (2019-nCoV) Real-Time RT-PCR assay. Cycle conditions were as follows: Standard setting; 50°C (10 min, 1 cycle), 95°C (2 min, 1 cycle), followed by 95°C (30s) and 55°C (3 s) for 45 cycles. Results were reported as genome copy number per 200 ng total RNA.

### Immunofluorescence

Tissues were prepared for immunostaining as previously described^52^. Briefly, viscera were fixed in 10% formalin and ganglia were fixed in 4% paraformaldehyde overnight, moved to 30% sucrose overnight, and subsequently embedded in optimal cutting temperature (OCT) media (ThermoFisher). A Leica CM3050-S cryostat (Leica Biosystems) was used to prepare 7 µm sections from each tissue block. Slides were rinsed in 1X PBS then blocked in 3% normal donkey serum, 0.1% Triton-100X, and 1X PBS for 30 min at room temperature. SARS-CoV-2 N protein was visualized using an Alexa Fluor® 488 conjugated rabbit monoclonal anti-SARS-CoV-2 nucleocapsid antibody at a 1:1000 concentration (NBP2-90988AF488; Novus Biologicals). SARS-CoV-2 spike protein was visualized using a mouse monoclonal anti-SARS-CoV-2 spike antibody at a 1:1000 concentration (GTX632604; GeneTex) followed by an Alexa Fluor®647 conjugated donkey anti-mouse polyclonal antibody at a 1:1000 concentration (ab150111; Abcam). dsRNA was visualized using a mouse monoclonal anti-dsRNA antibody at a 1:500 concentration (MABE1134-25UL; Sigma) followed by an Alexa Fluor®647 conjugated donkey anti-mouse polyclonal antibody at a 1:1000 concentration (ab150111; Abcam). hACE2 was visualized using a mouse monoclonal anti-ACE2 antibody at a 1:500 concentration (sc-390851; Santa Cruz Biotechnology) followed by an Alexa Fluor®647 conjugated donkey anti-mouse polyclonal antibody at a 1:1000 concentration (ab150111; Abcam). NeuN was visualized using an Alexa Fluor® 647 conjugated rabbit monoclonal anti-NeuN antibody at a 1:1000 concentration (ab190565; Abcam). α-d-galactose carbohydrate residues on sensory neurons was visualized using the *Bandeiraea simplicifolia* isolectin B4 (IB4) conjugated to rhodamine at a 1:250 concentration (RL-1102; Vector Laboratories). Tyrosine hydroxylase was visualized using an Alexa Fluor® 594 conjugated mouse monoclonal anti-TH antibody at a 1:500 concentration (818004; Biolegend). Glutamine synthetase was visualized using a mouse monoclonal anti-GS antibody at a 1:100 concentration (MA5-27750; Invitrogen) followed by an Alexa Fluor® 594 conjugated goat anti-mouse monoclonal antibody at a 1:1000 concentration (A11005; Invitrogen). Neuropilin-1 was visualized using a goat polyclonal anti-NRP-1 antibody at a 15 µg/mL concentration (AF566; R&D Systems) followed by an Alexa Fluor® 647 conjugated donkey anti-goat monoclonal antibody at a 1:1000 concentration (ab150135; Abcam). S100 beta was visualized using an Alexa Fluor®647 conjugated rabbit monoclonal anti-S100 beta antibody at a 1:1000 concentration (ab196175; Abcam). Iba1 was visualized using an Alexa Fluor® 647 conjugated rabbit monoclonal anti-Iba1 antibody at a 1:1000 concentration (ab225261; Abcam). Anti-phospho-Histone H2A.X (Ser139) was visualized using a mouse monoclonal anti-phospho-Histone H2A.X (Ser139) antibody at a 1:200 concentration (05-636; Millipore) followed by an Alexa Fluor®647 conjugated donkey anti-mouse polyclonal antibody at a 1:1000 concentration (ab150111; Abcam). Antibodies were validated in-house as described below. Nuclei were visualized with 4′,6-diamidino-2-phenylindole (DAPI) in SlowFade Diamond antifade mounting medium (ThermoFisher). Primary antibodies were incubated with tissues overnight at 4°C in 1% normal donkey serum, 0.1% Triton-100X, and 1X PBS. Secondary antibodies were incubated with tissues for 1 hour at room temperature. Mouse-on-mouse interference was reduced using Affinpure donkey polyclonal anti-mouse IgG Fab fragments at a 1:20 concentration (715-007-003; Jackson ImmunoResearch Laboratories Inc) for two hours preceding incubation with the unconjugated primary antibodies targeting dsRNA and SARS-CoV-2 spike as described above.

### Antibody validation

The anti-SARS-CoV-2 nucleocapsid antibody (NBP2-90988AF488) was validated by Novus Biologicals via western blotting. It was validated in our lab through immunostaining for SARS-CoV-2 nucleocapsid in SCGs and TGs (Fig. 1, Extended Data Fig. 2), DRGs and spine (Fig. 2, Extended Data Fig. 2,3), and brain (Fig. 3,4, Extended Data Fig. 3,4) from infected mice as well as uninfected SCGs, TGs, DRGs, spine, and brain from negative control mice. It was also validated via immunostaining of SCGs, TGs, DRGs, and brains from infected and uninfected negative control hamsters (Figure 7, Extended Data Fig. 7). It was additionally validated for *in vitro* use by staining infected and control primary neuronal cultures of SCGs, TGs, and DRGs (Fig. c,d,e, Extended Data Fig. 5,6). Immunostaining was assessed for punctate cytoplasmic staining of viral protein in infected tissue with absence in uninfected tissue. It was further validated in our lab by western blotting using homogenized SARS-CoV-2 WA1/2020 viral stock prepared for this study (Supplementary Fig. 1c). The blot was probed with the unconjugated parental antibody and an HRP conjugated goat polyclonal anti-rabbit IgG secondary antibody (ab6721; Abcam; Lot # GR34222167, Clone: polyclonal) and a band was observed just below 50 kDa, which is in line with the predicted molecular weight of 46 kDa, which is consistent with other published western blots for this protein^53–56^.Additional bands were noted between 37 kDa-25 kDa and 25 kDa-20 kDa, which is consistent with other validation western blots published by vendors for this protein as well as manuscripts listed above (Cell Signaling Technologies, Novus, Rockland, Proteintech). The anti-SARS-CoV-2 spike protein antibody (GTX632604) was validated by GeneTex by immunostaining, western blotting, ELISA, and immunoprecipitation. It was validated in our lab through immunostaining for SARS-CoV-2 spike in SCGs and TGs (Fig. 1, Extended Data Fig. 2), as well as DRGs (Fig. 2, Extended Data Fig. 2) from infected mice as well as uninfected SCGs, TGs, and DRGs from negative control mice. Immunostaining was assessed for punctate cytoplasmic staining of viral protein in infected tissue with absence in uninfected tissue. It was further validated in our lab by western blotting using homogenized SARS-CoV-2 WA1/2020 viral stock prepared for this study (Supplementary Fig. 1c). The blot was probed with the antibody and an HRP conjugated donkey polyclonal anti-mouse IgG secondary antibody (PA1-28664; Invitrogen; Lot # WI3375112, Clone: polyclonal) and a band was observed between 250-150 kDa, which is in line with the molecular weight of full-length spike (180-200 kDa), which is consistent with other published western blots for this protein^57–59^. The anti-dsRNA antibody (MABE1134-25UL), a marker of viral genome replication of RNA viruses, was validated by Sigma by immunostaining. It was validated in our lab through immunostaining for dsRNA in SCGs and TGs (Fig. 1, Extended Data Fig. 2), as well as DRGs (Fig. 2, Extended Data Fig. 2) from infected mice as well as uninfected SCGs, TGs, and DRGs from negative control mice. Immunostaining was assessed for punctate cytoplasmic staining of dsRNA in infected tissue with absence in uninfected tissue. As this antibody is specific for RNA, western blotting was not possible. The anti-hACE2 antibody (sc-390851), a cell membrane protein used by SARS-CoV-2 as a viral receptor, was validated by Santa Cruz via western blotting. It was validated in our lab through immunostaining of tissues from K18-hACE2 mice vs wild-type non-hACE2 expressing mice. Immunostaining was assessed for punctate membrane staining of hACE2 in tissue from K18-hACE2 mice with absence in wild-type non-hACE2 expressing mice (not shown). It was further validated in our lab by western blotting using a whole cell homogenate of HEK293 cells as a positive control (Supplementary Fig. 1c). It was further used to assess expression of hACE2 in whole tissue homogenate of SCGs, TGs, and DRGs from K18-hACE2 mice (Supplementary Fig. 1d). The blot was probed with the unconjugated parental antibody and an HRP conjugated donkey polyclonal anti-mouse IgG secondary antibody (PA1-28664; Invitrogen; Lot # WI3375112, Clone: polyclonal) and a band was observed between 100-150 kDa, which is in line with the predicted molecular weight of 100-110 kDa, which is consistent with other published western blots for this protein^60^. A mild elevation of ACE2 in the positive control (HEK293 cell homogenate) above 100 kDa is observed but is within the range of the molecular weight. It is not uncommon for ACE2 in HEK293 cell homogenate to appear mildly elevated above 100 kDa on western blots likely due to modifications in kidney epithelial cells that are not present in ganglia (R&D Systems). The anti-NeuN antibody (ab190565), a neuronal nuclear protein used as a marker of neurons, was validated by Abcam by western blotting of parental antibody. It was validated in our lab through immunostaining of neuronal tissues including SCGs and TGs (Fig. 1, Extended Data Fig. 2), DRGs and spine (Fig. 2, Extended Data Fig. 2,3), and brain (Fig. 3,4, Extended Data Fig. 3,4) from infected mice as well as uninfected SCGs, TGs, DRGs, spine, and brain from negative control mice. It was also validated via immunostaining of SCGs, TGs, DRGs, and brains from infected and uninfected negative control hamsters (Figure 7, Extended Data Fig. 7). Immunostaining was assessed for uniform nuclear staining with minimal cytoplasmic staining. It was further validated in our lab by western blotting using mouse whole brain homogenate (Supplementary Figure 1c). The blot was probed with the antibody and an HRP conjugated goat polyclonal anti-rabbit IgG secondary antibody (ab6721; Abcam; Lot # GR34222167, Clone: polyclonal) and doublet bands were observed flanking 50 kDa, which is in line with the molecular weight of 48 kDa, which is consistent with other published western blots for this protein^61–64^. The anti-neuropilin-1 antibody (AF566), a surface glycoprotein important in axon growth that binds VEGF in conjunction with tyrosine kinase receptors and is and putative SARS-CoV-2 receptor/co-receptor, was validated by R&D Systems by western blotting of knockdown samples. It was further validated in our lab by western blotting using a mouse whole brain homogenate as a positive control (Supplementary Fig. 1c). It was further used to demonstrate NRP-1 expression, via western blot, in whole tissue homogenate of SCGs, TGs, and DRGs from K18-hACE2 and wild-type mice (Fig. 8a, Supplementary Fig. 1c). The blots were probed with the antibody and an HRP conjugated donkey polyclonal anti-goat IgG secondary antibody (PA1-28664; Invitrogen; Lot # WI3375112, Clone: polyclonal) and a single band was observed between 100 kDa-75 kDa, which is in line with the reported molecular weight of 150-80 kDa, which is consistent with other published western blots for this protein^65–68^. Additional bands were noted between below 250 kDa and between 150 kDa-100 kDa, which is consistent with other validation western blots published by vendors for this protein as well as manuscripts listed above, which indicates recognition of both C and N-termini of the protein in positive control brain homogenate, which is to be expected as this antibody is polyclonal. (Cell Signaling Technologies, R&D, Proteintech, Abcam). Only the single banding pattern between 100 kDa-75 kDa was present in ganglia and is therefore used in the validation blot for the antibody and to verify presence in ganglia. The anti-tyrosine hydroxylase antibody (818004), a cytoplasmic marker of autonomic neurons, was validated by Biolegend by western blotting and by immunostaining of formalin-fixed paraffin-embedded tissues. It was previously validated in our lab through staining of positive control autonomic ganglia (SCG) vs negative control sensory ganglia (TG, DRG). Immunostaining was assessed for punctate cytoplasmic staining of TH in autonomic neurons with absence in sensory neurons. It was further validated in our lab by western blotting using mouse whole brain homogenate (Supplementary Fig. 1c). The blot was probed with the antibody and an HRP conjugated donkey polyclonal anti-mouse IgG secondary antibody (PA1-28664; Invitrogen; Lot # WI3375112, Clone: polyclonal) and doublet bands were observed between 75-50 kDa, which is in line with a molecular weight of 60 kDa, which is consistent with other published western blots for this protein^69–72^. The anti-glutamine synthetase antibody (MA5-27750), a cytoplasmic marker of satellite glial cells (SGCs), was validated by Invitrogen by knockout with western blotting. It was validated in our lab by immunostaining of SGCs in DRGs of infected mice (Figure 2g). It was validated for *in vitro* use by staining primary neuronal cultures of SCGs (Figure 5c, Extended Data Fig 5,6). Immunostaining was assessed for cytoplasmic staining of SGCs and absent staining of neurons. It was further validated in our lab by western blotting using mouse whole brain homogenate (Supplemental Figure 1c). The blot was probed with the antibody and an HRP conjugated donkey polyclonal anti-mouse IgG secondary antibody (PA1-28664; Invitrogen; Lot # WI3375112, Clone: polyclonal) and a band was observed between 50-37 kDa, which is in line with a calculated molecular weight of 42 kDa, which is consistent with other published western blots for this protein^73–76^. IB4 (RL-1102), a lectin that binds membrane bound α-d-galactose residues and is used as a marker of a sub-population of sensory neurons, was validated by Vector Laboratories by immunostaining positive control tissues. It was validated in our lab by immunostaining primary neuronal cultures of sensory TG and DRG neurons as well as autonomic SCG neurons (Fig. 5 c,d, Extended Data Fig. 5,6). Immunostaining was assessed for membrane staining of TGs and DRGs with absent staining of SCGs. Our immunostaining is similar to that which has previously been reported for this lectin^77–81^. As this is a lectin that binds a cell surface carbohydrate, western blotting was not possible. The anti-S100 beta antibody (ab196175), a cytoplasmic calcium binding protein used as a marker of astrocytes, was validated by Abcam by immunostaining. It was further validated in our lab by western blotting using mouse whole brain homogenate (Supplemental Figure 1c). The blot was probed with the antibody and an HRP conjugated goat polyclonal anti-rabbit IgG secondary antibody (ab6721; Abcam; Lot # GR34222167, Clone: polyclonal) and a band was observed just above 10 kDa, which is in line with a molecular weight of 11 kDa, which is consistent with other published western blots for this protein^82–84^. The anti-Iba1 antibody (ab225261), a cytoplasmic ionized calcium-binding adapter protein used as a marker of microglia, was validated by Abcam by immunostaining. The anti-phospho-Histone H2A.X (Ser139) antibody (05-636), a marker of phosphorylation of histone H2A.X at serine 139, which indicates DNA fragmentation associated with apoptosis, was validated by Millipore by immunoblotting. It was further validated in our lab by western blotting using whole cell homogenate of UV treated Vero E6 cells (Supplementary Fig. 1c). The blot was probed with the antibody and an HRP conjugated donkey polyclonal anti-mouse IgG secondary antibody (PA1-28664; Invitrogen; Lot # WI3375112, Clone: polyclonal) and a band was observed at between 20 kDa-15 kDa, which is in line with a molecular weight of 17 kDa, which is consistent with other published western blots for this protein^85, 86^.

### Confocal microscopy and image analysis

Imaging was performed using a Leica SP8 scanning confocal microscope. Sections of ganglia, brain, and spinal cord were imaged with identical laser power and gain settings within tissue type to account for background immunofluorescence. Cells from *in vitro* studies were imaged with varying laser power and/or gain due to the wide range of immunofluorescence observed within given experiments. Images were imported into ImageJ and contrast and brightness was adjusted identically across all images within tissue types. 3D models were made using ImageJ and SyGlass VR imaging software.

### Plaque assays

Flash frozen tissues were homogenized in Dulbecco’s Modified Eagle Medium (DMEM; Fisher Scientific) in bead tubes using a TissueLyser II (Qiagen) for 45 s sessions for three sessions. Ganglia were homogenized in 0.5 mL DMEM due to size. Brain regions and spinal cord segments were homogenized in 1.0 mL of DMEM. The undiluted tissue homogenate as well as a ten-fold dilution of homogenate was inoculated in duplicate onto confluent monolayers of Vero E6 cells in 24-well plates. After 1 h of adsorption, the inoculum was removed, a 0.5% agarose overlay added (DMEM with 8% fetal bovine serum, 1% penicillin/streptomycin, molecular grade agarose), and plates were returned to the incubator for 48 h at 37°C with 5% CO^2^. Plates were then fixed with 10% formaldehyde, the agarose overlay removed, and stained with plaque dye. Infectious viral titer is reported as plaque forming units per mL (PFU/mL) of tissue homogenate.

### Primary neuronal culture

Neurons from sensory ganglia (TG, LS-DRG) and autonomic ganglia (SCG) were collected from mature mice and cultured as previously described^87–95^. Briefly, ganglia were harvested from 8–10-week-old K18-hACE2 and WT mice and enzymatically digested with a sequential incubation of papain (Worthington) and collagenase/dispase (Sigma-Aldrich) followed by washes in Neurobasal A (Invitrogen) after each digestion. Ganglia were triturated into single cell suspensions via pipette and further washed. SCGs and LS-DRGs were brought to volume in “complete media” containing Neurobasal A with 2% B27 (Invitrogen), 1% penicillin-streptomycin (Thermo Fisher), Glutamax (Thermo Fisher), fluorodeoxyuridine (Sigma-Aldrich), and neurotrophic factors (PeproTech). Neurons were plated at a concentration of 3,000 neurons/well in Matrigel coated Lab-Tek II 8-well chamber slides or 24-well plates (Thermo Fisher). TGs were collected after separation from debris via density gradient centrifugation using OptiPrep (Sigma-Aldrich) with subsequent washes in Neurobasal A. TGs were plated as described for SCGs and LS-DRGs. For *in vitro* neuronal culture studies mice were genotyped using the Jackson Laboratory assay for verification of transgene expression using cDNA from a sample from each pooled ganglia taken before plating into replicates from each neuronal culture session (Supplementary Fig. 1b). Neuronal cultures were allowed to recover for 3 days *ex vivo* prior to infection.

### Neuronal infection

Neurons from K18-hACE2 and WT mice were inoculated with SARS-CoV-2 isolate USA-WA1/2020 at 30 MOI in 100 µL Neurobasal A (Invitrogen) for 8-well chamber slides and 200 µL for 24-well plates for 1 h. Following the 1 h adsorption the inoculum was removed, fresh complete media was added (without fluorodeoxyuridine), and neurons were incubated at 37°C with 5% CO2. Aliquots of the inocula and viral stock were saved for back titration using plaque assays and RT-qPCR.

### Quantification of infection in autonomic and sensory ganglia in primary neuronal culture and tissues

To quantify the number of autonomic (SCG) and sensory (DRG) neurons infected per ganglia *in vitro*, 8-well chamber slides containing SCGs and DRGs from hACE2 and WT mice were fixed with paraformaldehyde at 1-, 2-, and 3-dpi and stained as described above for the detection of SARS-CoV-2 nucleocapsid. The number of infected neurons from each ganglion were counted for each day, averaged, and reported as the percentage of infected neurons per 500 neurons counted. LS-DRG neurons were chosen as the representative sensory neuron as they had the more dynamic replication kinetics with successive rounds of replication. To quantify the number of autonomic (SCG) and sensory (LS-DRG, TG) neurons infected per ganglia *in vivo*, tissue sections from hACE2 and WT mice were immunostained as described above for SARS-CoV-2. The number of infected neurons from each ganglion were counted across three tissue sections spanning the ganglion being counted, averaged, and reported as described above.

### Detection of SARS-CoV-2 assembly and release from neurons

To determine if neurons from K18-hACE2 and WT mice are permissive to infection and release of infectious virus, neurons were incubated for up to 5 dpi (depending on availability of neurons from the specific ganglia) with daily sampling. To determine the amount of virus bound to neurons vs that left unbound immediately following incubation with the inoculum, the inoculum and neurons were collected separately and constitute the 0-dpi sample. Daily, media and neurons were collected separately in duplicate (TGs, LS-DRGs) or singularly (SCGs) in 500 µL of LS-TRI Reagent (Fisher Scientific) for RNA extraction and viral genome copy number quantitation via RT-qPCR as described above. Samples were stored at 4°C until processing. For quantification of viral titer in neurons vs that released into the media, neurons and media were collected separately in duplicate (TGs, LS-DRGs) or singularly (SCGs). To correct for evaporation of media throughout the time course the final volume of collected media was brought up 500 µL by adding DMEM prior to plaque assay. Neurons were collected in 500 µL DMEM after scraping with a pipette tip. Samples were immediately stored at −80°C until processing for plaque assay as described above. Following collection of the media but prior to collection of the neurons in TRI reagent or DMEM, the neurons were gently washed with 500 µL DMEM which was then discarded, to remove any residual media containing RNA or virus. A similar rinse was performed immediately after the 1 h inoculation to remove any residual inoculum.

### Inhibition of SARS-CoV-2 infection by neuropilin-1 blockade in primary sensory neuronal culture using a small molecule inhibitor or blocking antibody

Primary neuronal cultures of LS-DRGs from K18-hACE2 and WT mice were established as described above. Neurons were pretreated with 100 µM of the NRP-1 antagonist EG00229 (6986; Tocris) dissolved in DMSO prior to infection as described for Caco-2 cells^26^. EG00229 putatively blocks binding between the carboxyl-terminal sequence of SARS-CoV-2 S1 which has a C-end rule (CendR) motif and the extracellular b1b2 CendR binding pocket of NRP-1, which has been suggested as an alternative co-receptor for SARS-CoV-2 in non-neuronal cells ^26–28^. Neurons were infected as described above. To determine if NRP-1 blockade impacted SARS-CoV-2 entry and therefore subsequent replication in neurons, neurons and media were collected together in LS-TRI Reagent (Fisher Scientific). RNA was isolated and virus replication assessed via RT-qPCR as described above. Samples were collected at initial peak replication times as determined through our previous neuronal growth kinetics studies (LS-DRG; 3 dpi) to assess if theses peaks were blunted or completely inhibited. Infected neurons from K18-hACE2 and WT mice not treated with EG00229 but with an equivalent amount of DMSO, the solvent for EG00229, served as controls. Neuropili-1 inhibition studies were repeated as described above for EG00229 using a goat polyclonal anti-NRP-1 blocking antibody (AF566; R&D Systems) at 1 µg/mL, 15 µg/mL, and 30 µg/mL. This antibody was shown by the vendor to block 50% of binding of the NRP-1 ligand VEGF at 0.3-1 µg/mL and >90% of binding at 30 µg/mL.

### Detection of hACE2 and NRP-1 in PNS ganglia

Expression of hACE2 in PNS tissues of K18-hACE2 mice as well as NRP-1 expression in K18-hACE2 and WT mice was assessed by standard western blotting (Supplementary Fig. 1d,e). In summary, ganglia from SCGs, TGs, DRGs were collected from each mouse type and homogenized in RIPA buffer, protein concentration determined via Bradford assay (BioRad), and 15 µg of total protein loaded into the stacking gel and subjected to SDS-PAGE containing trichloroethanol. HEK293 cell homogenate served as a positive control for hACE2. A mouse whole brain homogenate served as a control for NRP-1. Proteins were transferred to a PVDF membrane using a wet transfer. Blots were imaged for total protein using activation of trichloroethanol in a BioRad Chemidoc gel imager (BioRad). Blots were blocked overnight in a cold room in a solution of 5% milk in TBST with Tween. Primary antibody for hACE2 or NRP-1 as listed above were added to blots and incubated for four hours in a cold room. Blots were washed and then probed with appropriate HRP conjugated secondary antibodies as listed above for 1 hour. Blots were washed and imaged on a BioRad Chemidoc gel imager after applying SuperSignal West Femto Maximum Sensitivity Substrate (Thermo).

### Statistics and reproducibility

Sample sizes were not statistically calculated as they are similar to sample sizes used in other SARS-CoV-2 studies using K18-hACE2 mice^14, 15, 19, 21^ or golden Syrian hamsters^96–98^. Animals were randomly assigned to either inoculum group or control group ensuring the groups were age-matched. Measurements were taken from distinct samples. RT-qPCR and plaque assays were performed in duplicate for each sample when assessing both *in vivo* and *ex vivo* infections. RT-qPCR results that fell below the lower limit for the standard curve (8 copies) after normalization were reported as 0 for inclusion in analysis, no data were excluded. Neuronal infection studies, both immunostaining as well as plaque assay and RT-qPCR studies, were repeated in three separate experiments, with duplicate samples for each ganglion and timepoint in K18-hACE2 and in duplicate in WT mice. Neuropilin-1 inhibition studies were repeated twice, with duplicate samples for each timepoint in K18-hACE2 and WT mice. Mouse infection studies were repeated as described. Golden Syrian hamster infection studies were conducted as described. All statistical analyses were performed in JMP Pro 16 (SAS Institute) and confirmed in GraphPad Prism version 8 during figure creation. For statistical analysis, significance was set at *p* < 0.05, calculated as two-tailed. RT-qPCR data was log transformed before analysis to correct for normality of distribution. RT-qPCR data was analyzed using a multifactorial ANOVA. If significance was found, pairwise analysis was performed using Tukey’s honestly significant difference (HSD) post hoc test. Inhibitor studies were analyzed using unpaired two-tailed t-tests.

### Data availability

Data generated during this study and referenced in this manuscript are available from the corresponding author upon reasonable request.

## Acknowledgments

This research was funded by internal COVID-19 rapid response seed funding from the Fralin Life Sciences Institute at Virginia Tech (ASB). The following reagent was deposited by the Centers for Disease Control and Prevention and obtained through BEI Resources, NIAID, NIH: SARS-Related Coronavirus 2, Isolate USA-WA1/2020, NR-52281. Special thanks to Jonathan Auguste, Will Stone, Addie Hayes, and Michael Bowers for various forms of assistance.

## Author contributions

Conceptualization, JDJ, ASB; methodology, JDJ, NKD, CKT, ASB; validation, JDJ, CKT, ASB; formal analysis, JDJ, GAM, CKT, ASB; investigation, JDJ, GAM, PG, TLH, TMT, SAH, JCG, MJ, NY, EHL, CKT, ASB; resources, NKD, CKT, ASB; data curation, JDJ, CKT, ASB; writing—original draft preparation, JDJ; writing—review and editing, JDJ, NKD, CKT, ASB; visualization, JDJ, CKT, ASB; supervision, NKD, CKT, ASB; project administration, ASB; funding acquisition, ASB. All authors have read and agreed to the published version of the manuscript.

---

**Extended Data Fig 1.**
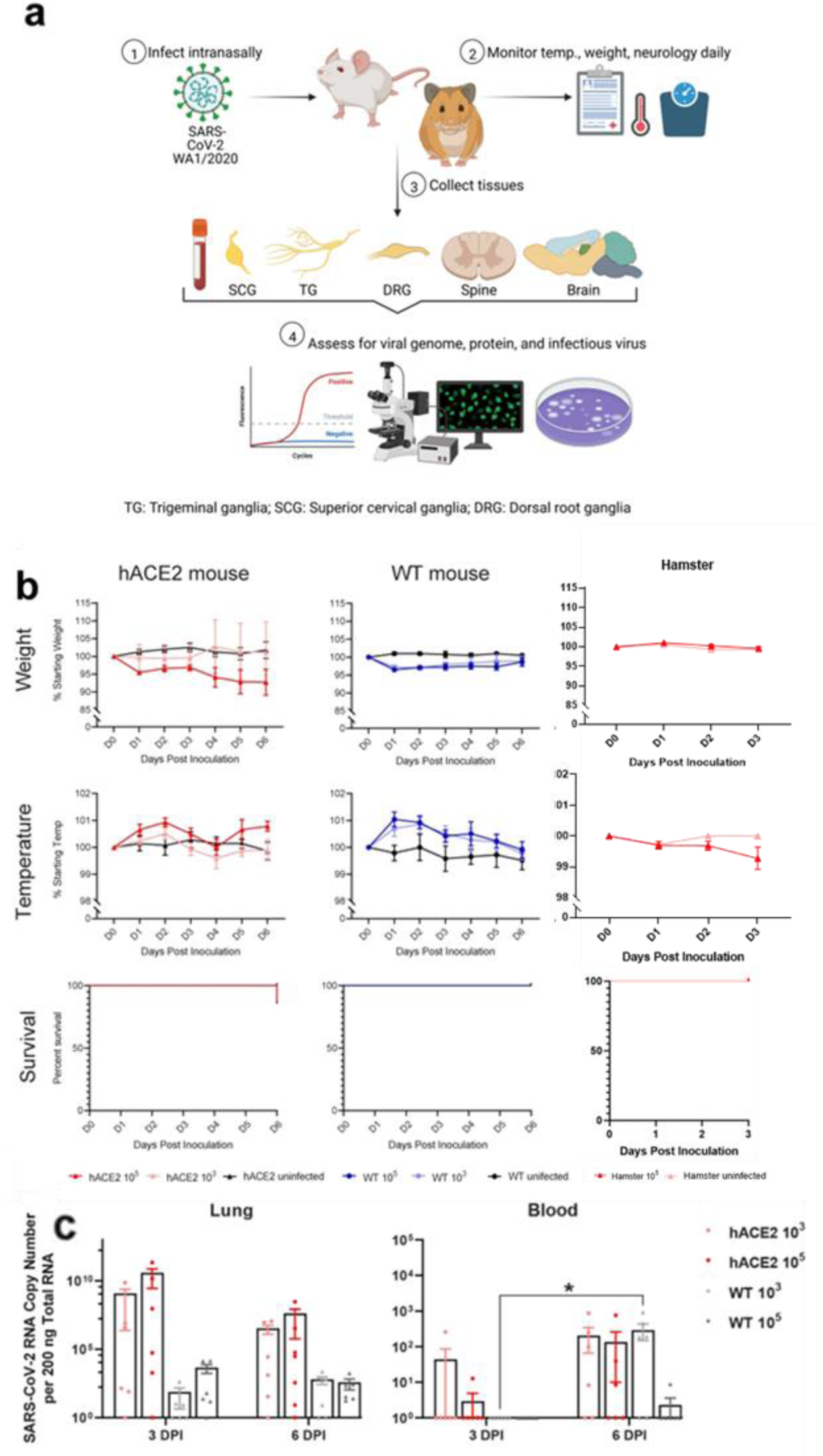
Experimental approach and clinical data for hACE2 mice, WT mice, and golden Syrian hamsters. **a**, Graphical abstract outlining the experimental approach used in mouse (hACE2: n=35; WT: n=34; controls: n=4/mouse type) and Syrian golden hamster (n=9; control: n=1) infections highlighting intranasal infection of groups with either 3 log PFU or 5 log PFU SARS-CoV-2 (mice) or 5 log PFU SARS-CoV-2 (hamsters), clinical evaluation (temperature, weight, survival, von Frey threshold), collection of tissues, and downstream analysis for SARS-CoV-2 RNA copies (RT-qPCR), virus and host antigen (immunostaining), and infectious virus (plaque assay). **b**, Clinical data by inoculum group for mice and hamsters including weight, temperature, and survival. Weight (grams) for each animal was recorded daily and reported as the mean percentage increase/decrease (±SD) for each inoculum group relative to the mean starting weight for that group. The only group to exhibit notable weight loss was the hACE2 mice inoculated with 5 log PFU, which began to lose weight after 3 dpi. A minor but insignificant decrease in weight was observed in infected hamsters relative to uninfected control. Temperature (°C) for each animal was recorded daily. Temperature is reported as the mean percentage increase/decrease for each inoculum group relative to the mean starting temperature for that group. A transient temperature increases occurred in the 5 log PFU inoculated mice whereas a mild decrease occurred in 5 log PFU inoculated hamsters. Kaplan-Meier survival plots were created for each inoculum group. The only group to have mortality was the hACE2 mice inoculated with 5 log PFU. Mortality was noted at 6 dpi (14%, n= 2 of 14). **c**, SARS-CoV-2 RNA was detected in lungs of hACE2 and WT mice in both inocula groups at both timepoints, which decreased over time. While differences were detected in the lungs (F(7, 41) = 2.745, p = 0.0197) none were between relevant groups. Low concentrations of SARS-CoV-2 RNA were detected in the blood of hACE2 in both inoculum groups at 3 dpi and both hACE2 and WT mice in both inoculum groups at 6 dpi. A significant difference (F(7, 40) = 3.417, P = <0.006) was detected in the WT group inoculated with 10^3^ PFU assessed at 3- vs 6-dpi (p= 0.0321). Data are the mean ± s.e.m. Log transformed RNA genome copy numbers were statistically compared by three-way ANOVA (independent variables: inocula, days post infection, genotype). Pairwise comparisons were conducted using Tukey’s HSD post hoc tests. *p < 0.05, **p < 0.01, ***p < 0.001. See Supplementary Figure 1 for hACE2 genotyping, hACE2 protein expression, and RT-qPCR controls.

**Extended Data Fig 2.**
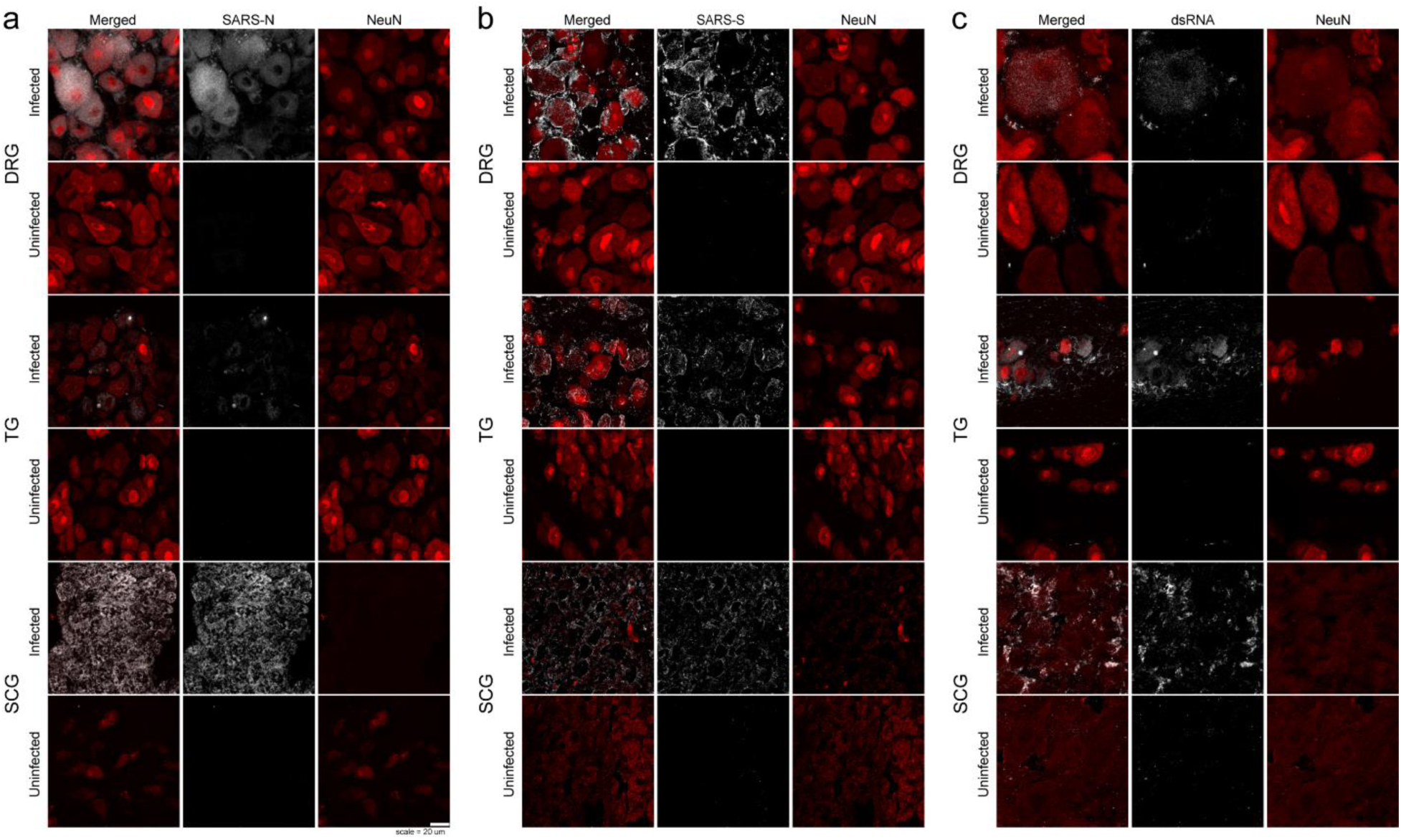
Immunofluorescence for SARS-N protein, SARS-S protein, dsRNA, and NeuN in peripheral ganglia from hACE2 mice. All images were acquired using a Leica SP8 confocal microscope, using identical image acquisition settings (laser power and gain) across all sections shown for each SARS-related antibody. All images were colorized, z-projected, and prepared using identical contrast and brightness parameters in ImageJ for each SARS-related antibody. a, SARS-N is present in neurons in the LS-DRG, TG and SCG in infected mice. Minimal background immunofluorescence is observed in sections from uninfected mice. Of note is substantial vacuolization and loss of NeuN immunofluorescence in the infected SCG. b, SARS-S is present in neurons in the LS-DRG, TG and SCG in infected mice, whereas there is some extracellular nonspecific staining visible, likely due to mouse-on-mouse immunohistochemistry artifacts. c, dsRNA is present in neurons and some satellite glial cells in the LS-DRG, TG and SCG in infected mice. Similar to SARS-S, some nonspecific immunostaining is present, also likely due to mouse-on-mouse immunostaining artifacts. The difference between infected and uninfected cells is readily apparent, however, for SARS-S and dsRNA. See Supplementary Figure 1 for additional antibody validation via western blot, hACE2 genotyping, and hACE2 protein expression.

**Extended Data Fig 3.**
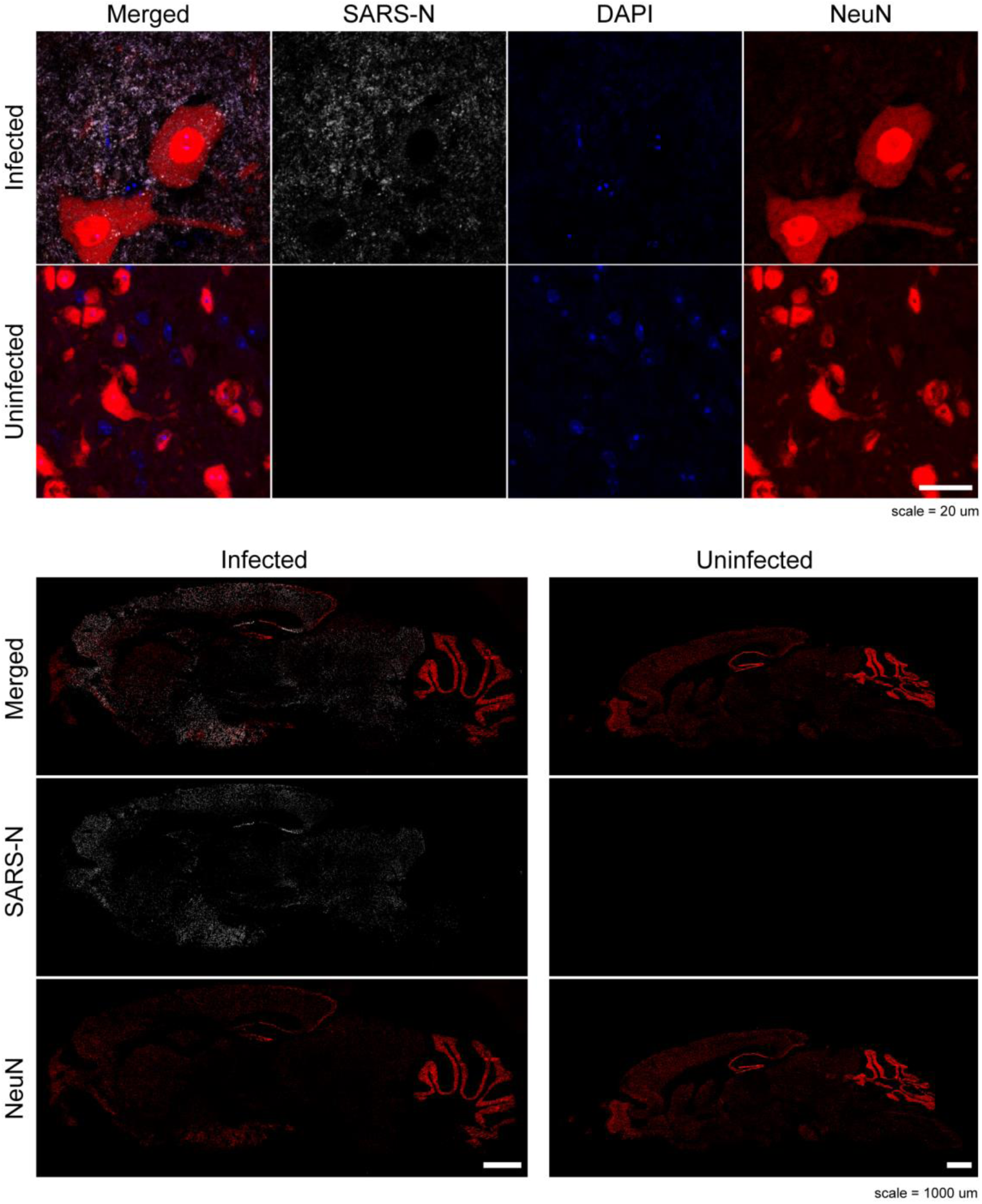
Immunofluorescence for SARS-N protein and NeuN in sections of spinal cords and brains taken from infected and uninfected hACE2 mice. All images were acquired using a Leica SP8 confocal microscope, using identical image acquisition settings (laser power and gain) across all sections from each tissue type shown. All images were colorized, z-projected, and prepared using identical contrast and brightness parameters in ImageJ. SARS-N is present in neurons in the spinal cord and brains in infected mice. Minimal background immunofluorescence is observed in sections from uninfected mice. Of note, SARS-N intensity was qualitatively substantially higher in the brain than in the spinal cord. See Supplementary Figure 1 for additional antibody validation via western blot, hACE2 genotyping, and hACE2 protein expression.

**Extended Data Fig 4.**
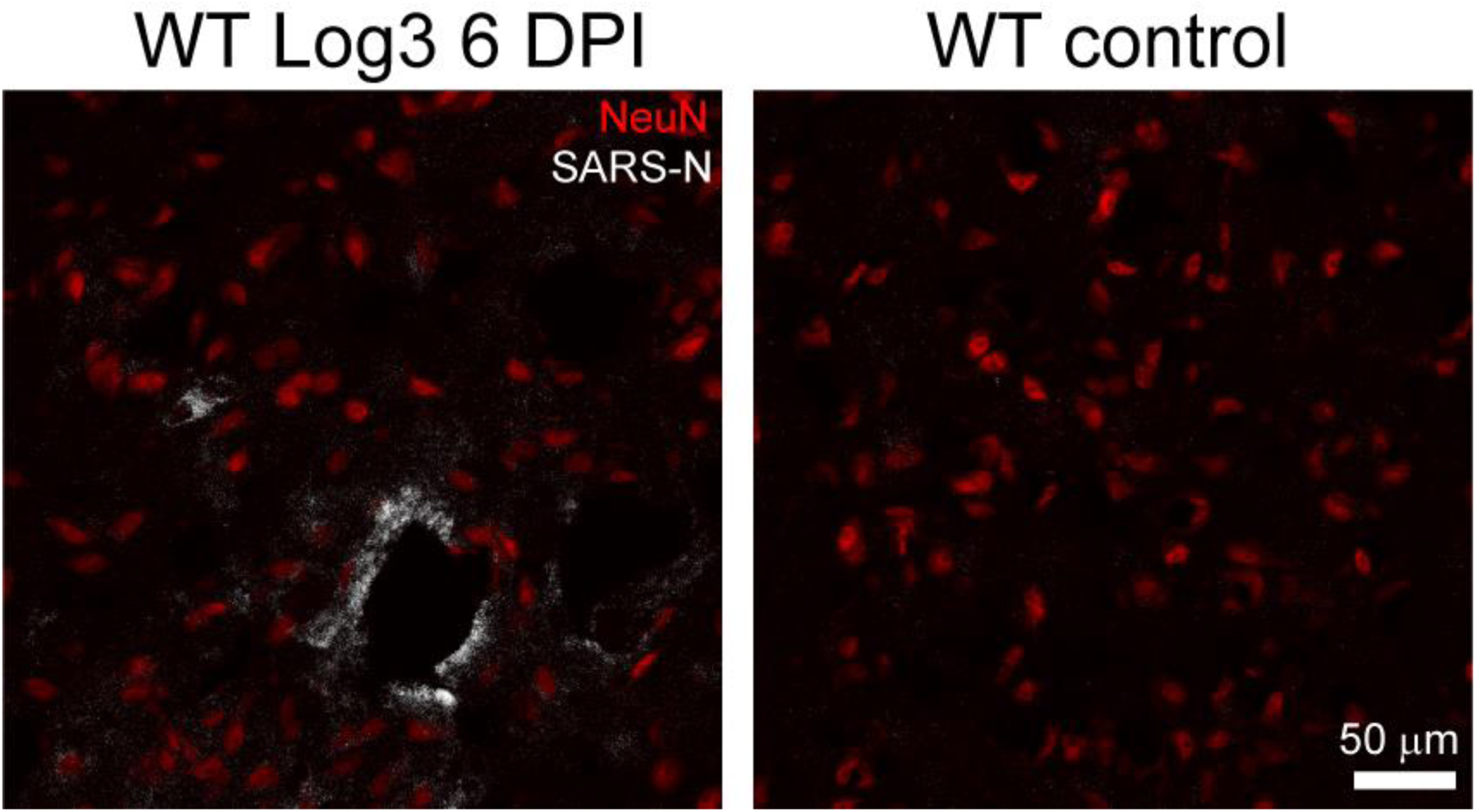
Immunofluorescence for SARS-N protein and NeuN in sections of brains taken from infected and uninfected WT mice. All images were acquired using a Leica SP8 confocal microscope, using identical image acquisition settings (laser power and gain) across all sections from each tissue type shown. All images were colorized, z-projected, and prepared using identical contrast and brightness parameters in ImageJ. SARS-N is present in a minority of neurons in infected WT mice. Of note, heavy SARS-N staining can be seen in a circular ring of tissue, possibly vasculature, with scattered SARS-N signal throughout the surrounding area. Minimal background immunofluorescence is observed in sections from uninfected mice. It is worth noting that while SARS-N was minimally detected by immunostaining in the brains of WT mice, as reported by other groups, SARS-CoV-2 RNA was readily detected by RT-qPCR in discrete brain regions at 3 and 6 dpi. Also, infectious virus was recovered via plaque assay from the hippocampi (3 log group: 2 PFU/mg homogenate; 5 log group: 5 PFU/mg homogenate) and brainstem (3 PFU/mg homogenate) of WT mice at 6 dpi. The use of multiple complimentary assays when applied to discrete brain regions increases the likelihood of detecting neuroinvasion in these mice. See Supplementary Figure 1 for additional antibody validation via western blot.

**Extended Data Fig 5.**
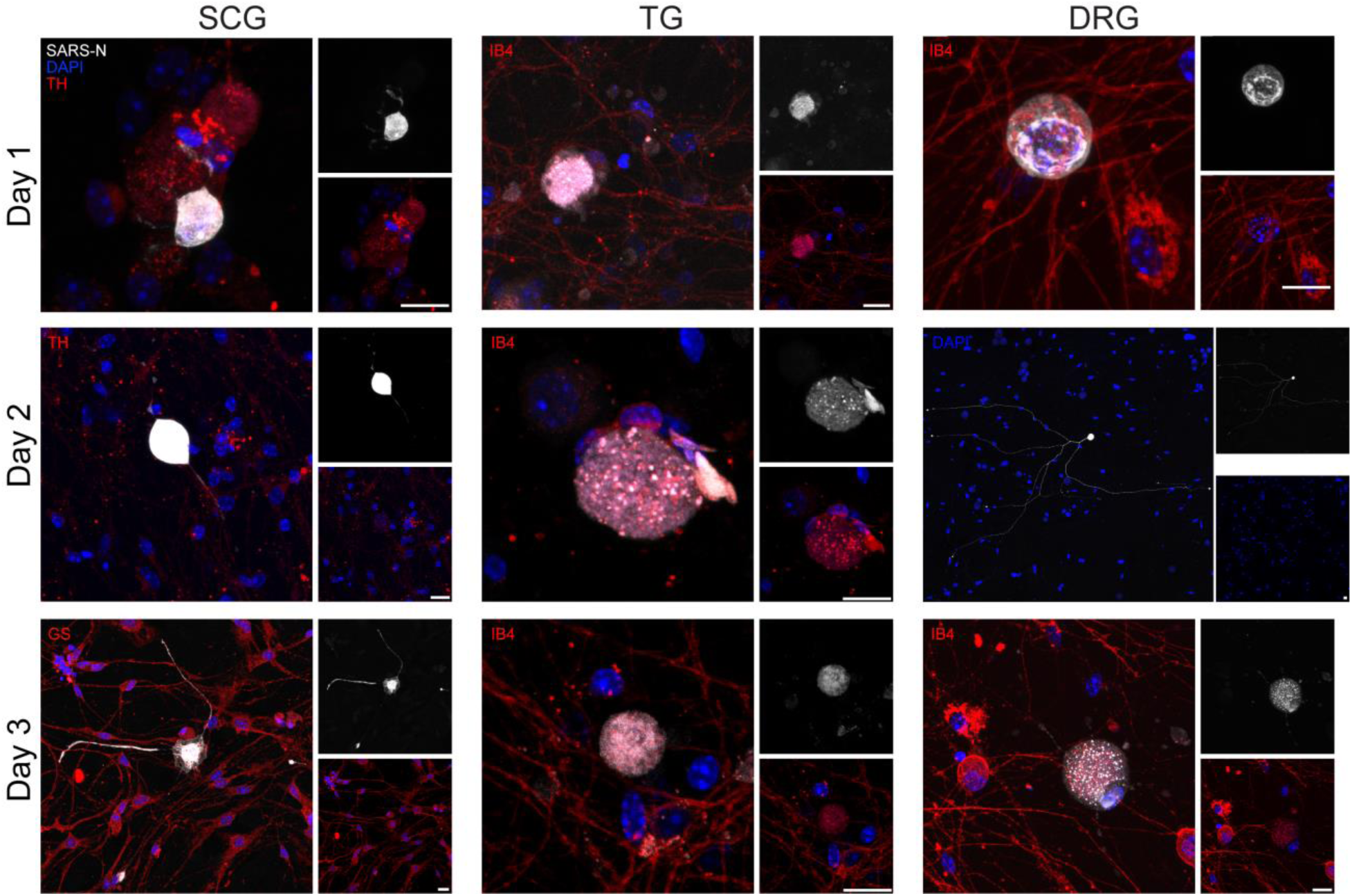
Immunofluorescence for SARS-N protein in *in vitro* cells taken from hACE2 peripheral ganglion cultures, incubated for two days, then inoculated with SARS-CoV-2. Cells were fixed 1-3 dpi and stained for SARS-N and various counterstains. All images were acquired using a Leica SP8 confocal microscope. Because of substantial variability in intensity of SARS-N immunofluorescence, laser power and gain were adjusted in order to highlight features of each cell. Day 2 LS-DRG is a montage image to show SARS-N detected throughout the neurites of one infected neuron; only DAPI is shown. SARS-N is present in neurons in the SCG, TG, and LS-DRG in infected mice. TH = tyrosine hydroxylase, IB4 = Isolectin-B4, GS = glutamine synthetase, SARS-N = SARS-CoV-2 nucleocapsid, DAPI = 4′,6-diamidino-2-phenylindole. Scale bar = 20 μm. See Supplementary Figure 1 for additional antibody validation via western blot, hACE2 genotyping, and hACE2 protein expression. See Supplementary Figure 6 for uninfected neuronal culture control images.

**Extended Data Fig 6.**
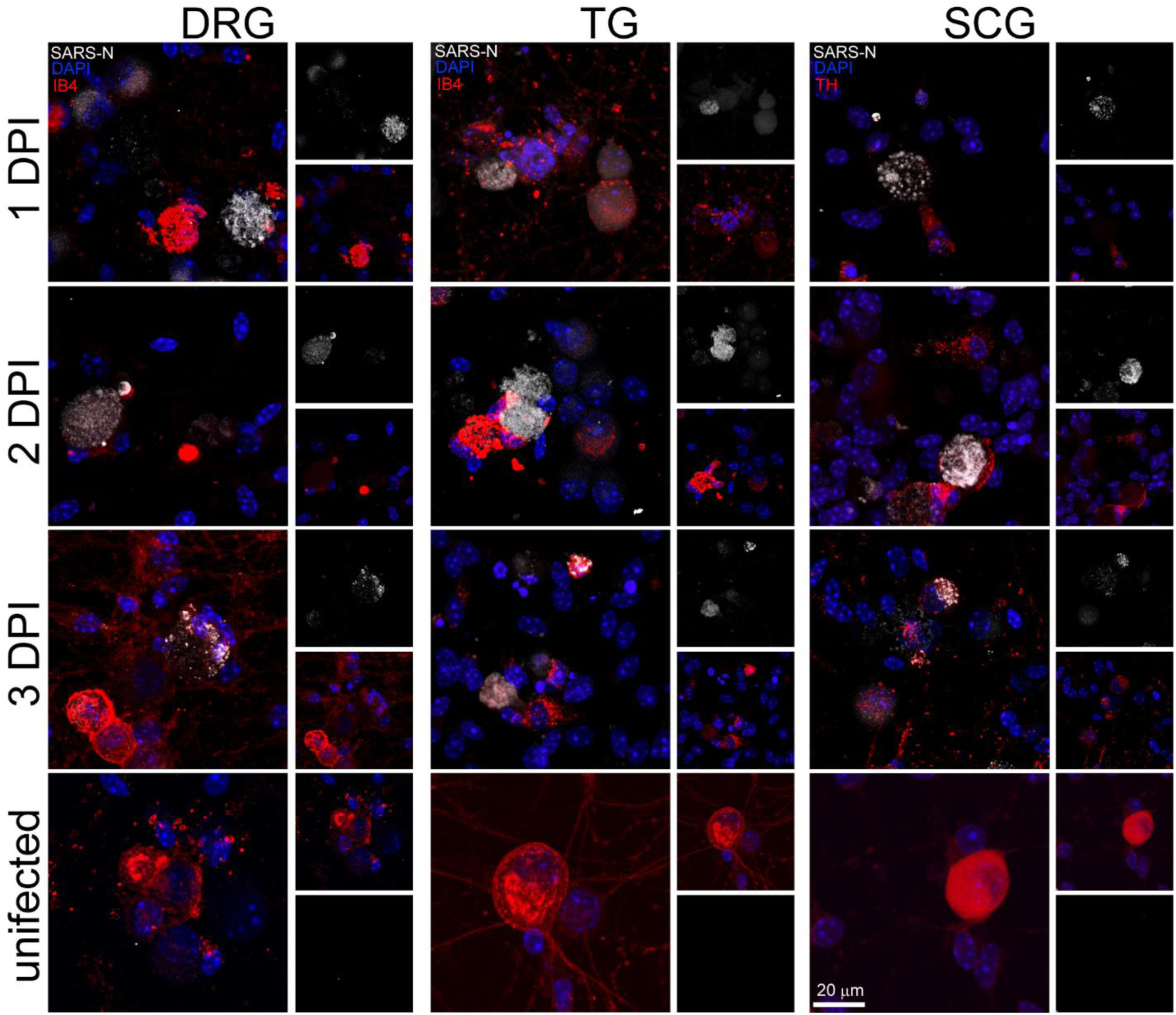
Immunofluorescence for SARS-N protein in *in vitro* cells taken from WT peripheral ganglion cultures, incubated for two days, then inoculated with SARS-CoV-2. Cells were fixed 1-3 dpi and stained for SARS-N and various counterstains. All images were acquired using a Leica SP8 confocal microscope. Punctate SARS-N immunoreactivity is visible in many infected cells. Virtually no background is visible in uninfected cells. Identical imaging and post-processing parameters were used for all images. TH = tyrosine hydroxylase, IB4 = Isolectin-B4, GS = glutamine synthetase, SARS-N = SARS-CoV-2 nucleocapsid, DAPI = 4′,6-diamidino-2-phenylindole. Scale bar = 20 μm. See Supplementary Figure 1 for additional antibody validation via western blot.

**Extended Data Fig 7.**
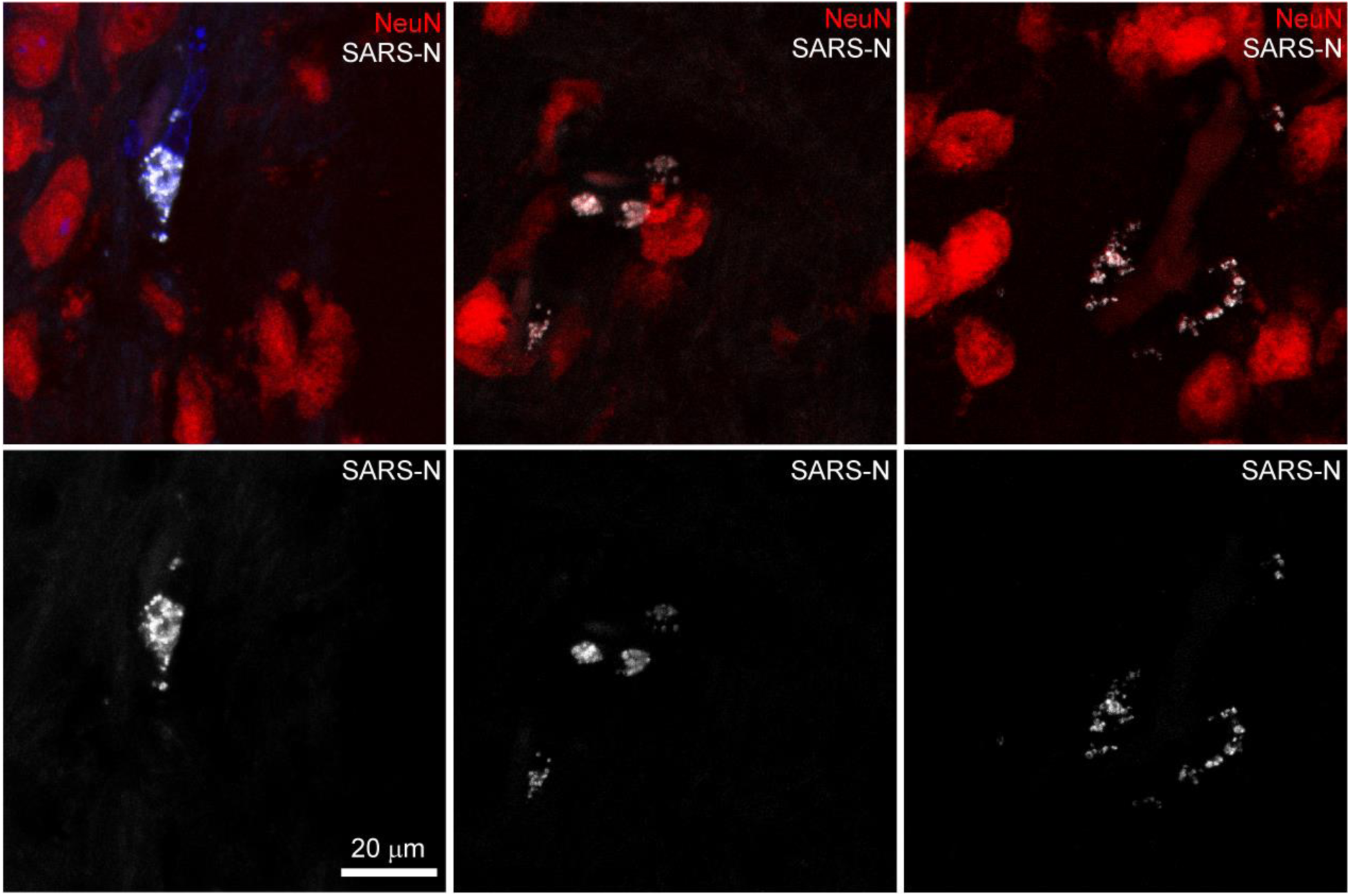
Immunofluorescence for SARS-N protein and NeuN in sections of brains taken from infected golden Syrian hamsters. All images were acquired using a Leica SP8 confocal microscope, using identical image acquisition settings (laser power and gain) across all sections from each tissue type shown. All images were colorized, z-projected, and prepared using identical contrast and brightness parameters in ImageJ. SARS-N is present in some cells in infected hamster brains, which appear to be associated with vasculature. See Supplementary Video 5 for 3D rendering of this image. See Supplementary Figure 1 for additional antibody validation via western blot.

**Supplementary Fig 1.**
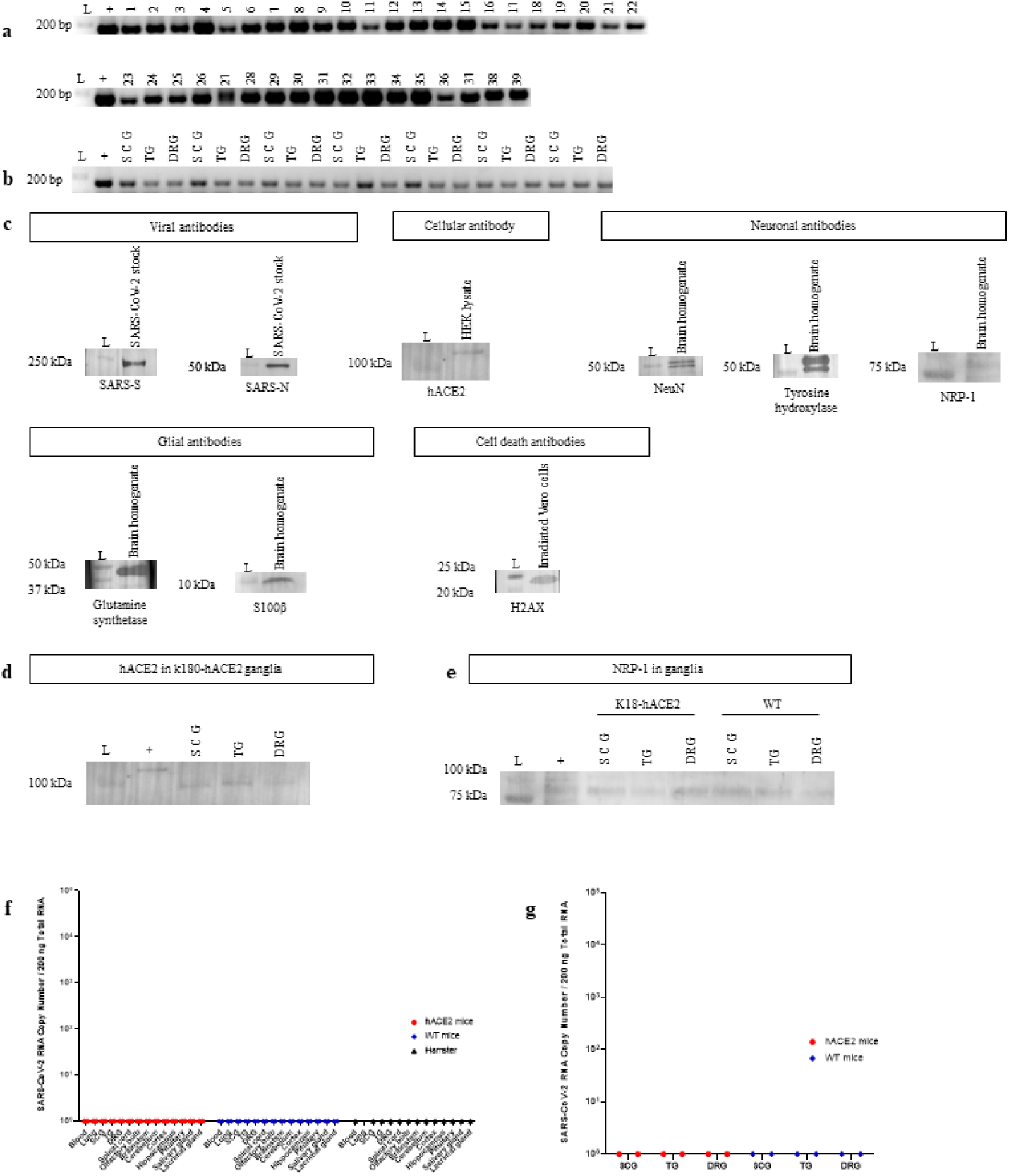
Validation of hACE2 mice, immunostaining antibodies, and SARS-CoV-2 N1 specific RT-qPCR assay. **a**, hACE2 mice (n=39; 35 infected, 4 controls) were genotyped following Jackson Laboratory protocol #38170 V2. The presence of a PCR product at ≈170 bp, the predicted size of the hACE2 amplicon, confirms presence of the transgene in these mice. **b**, SCGs, TGs, LS-DRGs used for neuronal culture infection studies (n=7 independent cultures) were genotyped as above to confirm presence of the transgene. The presence of an amplicon at ≈170 bp confirms presence of the transgene in these primary neuronal cultures. **c**, Antibodies used in immunostaining were validated by western blot using positive controls recommended by their manufacturers. **d**, Expression of hACE2 protein was confirmed via western blot in homogenates of SCGs, TGs, and LS-DRGs from hACE2 mice by observation of a band at ≈100 kDa (molecular weight 100-110 kDa). A mild elevation of ACE2 in the positive control (HEK293 cell homogenate) above 100 kDa is observed but is within the range of the molecular weight. It is not uncommon for ACE2 in HEK293 cell homogenate to appear mildly elevated above 100 kDa on western blots likely due to modifications in kidney epithelial cells that are not present in ganglia (R&D Systems). **e**, Expression for NRP-1 protein was confirmed via western blot in homogenates of SCGs, TGs, and LS-DRGs from hACE2 and WT mice. **f**, The SARS-CoV-2 N1 specific RT-qPCR assay was validated for each tissue assessed in hACE2 mice, WT mice, and hamsters by assessing negative control tissues from uninfected hACE2 mice (n=2), WT mice (n=2), and hamsters (n=1). **g**, The SARS-CoV-2 N1 specific RT-qPCR assay was validated for neuronal culture infection studies by assessing negative control uninfected cultures of SCGs, TGs, and LS-DRGs from hACE2 and WT mice (n=2 per ganglia per mouse type). Details on genotyping, immunostaining validation, and western blotting are described in the methods section and in the Reporting Summary.

**Supplementary Fig 2.**
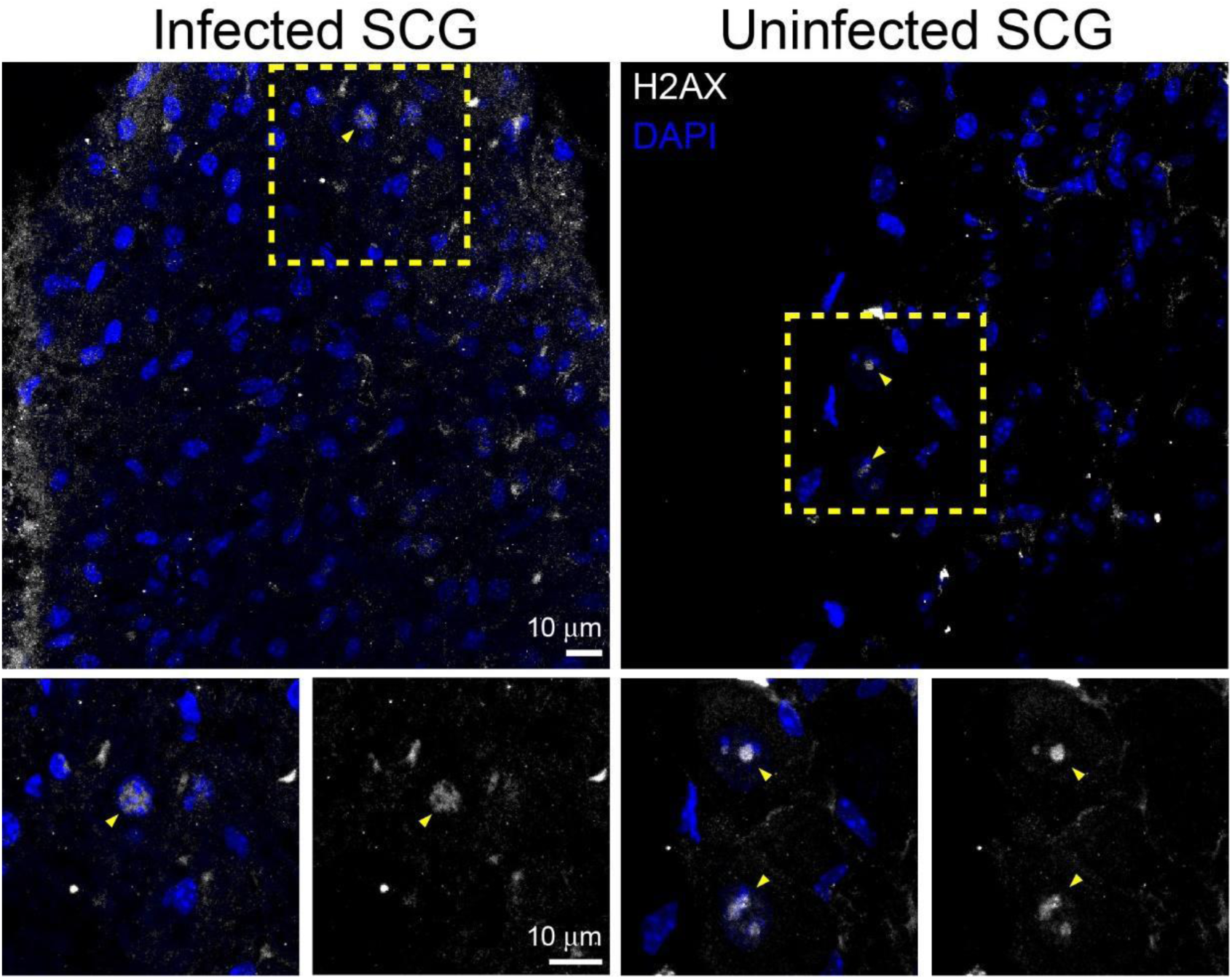
Immunofluorescence for phospho-histone H2A.X in the SCG of hACE2 mice. Given the apparent pathology noted in the SCGs of hACE2 and WT mice, the SCG was immunostained for H2A.X, a marker of DNA fragmentation and apoptosis. H2A.X immunoreactivity was observed in only a few SCG cells in both infected and uninfected tissues, indicating that SARS-CoV2-induced pathology in SCG did not result in extensive DNA damage, at least at the time points examined, suggesting other mechanisms of cell cytopathology. See Supplementary Figure 1 for additional antibody validation via western blot, hACE2 genotyping, and hACE2 protein expression.

